# A conserved paint box underlies color pattern diversity in Estrildid finches

**DOI:** 10.1101/2021.02.19.431992

**Authors:** Magdalena Hidalgo, Camille Curantz, Nicole Quenech’Du, Thanh-Lan Gluckman, Julia Neguer, Samantha Beck, Ammara Mohammad, Marie Manceau

## Abstract

Many animals exhibit typical color patterns that have been linked to key adaptive functions, yet the developmental mechanisms establishing these crucial designs remain unclear. Here, we surveyed color distribution in the plumage across a large number of passerine finches. Despite extreme apparent pattern diversity, we identified a small set of conserved color regions whose combinatory association can explain all observed patterns. We found these domains are instructed by early embryonic landmarks, and through profiling and comparative analyses produced a molecular map marking putative color domains in the developing skin. This revealed cryptic pre-patterning common to differently colored species, uncovering a simple molecular landscape underlying extensive color pattern variation.

---

Systematics ever refines the description of phenotypic variation in vertebrates, shedding light on a fascinating paradox: morphological diversity is extensive, associated to a breadth of behavioral and physiological function; yet, it is bound to an underlying order, as within taxa most color patterns and shapes display conserved orientation, geometry, or periodicity (1–3). Such trends are likely rooted in the spatio-temporal hierarchy of genetically determined developmental programs, unfolding to progressively refine space in the embryo from the early establishment of body axes to the precise positioning of tissue domains (e.g., brain regions, limb segments (4,5)). This mechanism however hardly reconciles with extreme and/or rapid phenotypic change. Conversely, spontaneous spatial arrangement through tissue “selforganization” may reach stable states corresponding to complex morphologies (e.g. periodic coloration, teeth circumvolutions) that vary importantly through minimal parameter change (6,7). This inherent malleability provides an appealing explanation to morphological diversity, but does not ensure reproducibility. For lack of comprehensive work uncovering the developmental origin(s) of extensive phenotypic variation in a vertebrate system, the mechanisms producing adaptive differences in morphologies while guaranteeing functional stability remain unknown.

Here, we took advantage of color pattern, trait crucial for fitness, and its broad variation in finches of the *Estrildidae* family of passerine songbirds. In this monophyletic group, sexual signal driven plumage coloration ranges from homogenous to compartmented in large areas or in periodically repeated geometries (8). We described the distribution of pigments in the adult plumage on dorsal and ventral flat skin preparations of 38 species, covering most frequently encountered genus in the family (Supplementary Figure 1, (9)). We characterized feather types according to distal hue and motif, all feathers having a proximal grey basis (Supplementary Figure 2). We focused on breast and trunk regions of feather implantation (i.e., tracts) where contrary to the head feather follicles are arranged in chevrons, typical structures whose number and feather arrangement is highly conserved, providing spatial reference along both body axes (Figure 1A, (10)). Recording feather types at each position in dorsal and ventral tracts showed they never intermingle –though they may form gradients, creating well-defined color domains. While some boundaries extended over a few chevrons, most were sharp, and all had highly reproducible mean location independent of gender and age (Figure 1B for the ventrum, Supplementary Figure 3 for the dorsum, and Supplementary Figure 4 for methods). Strikingly, we observed recurring overall boundary locations (e.g., in 25 species, ventral bilateral sides are split in a flank and a ventral domain) and orientation, most being parallel to tract axes (Supplementary Figure 5). We built reference maps of most frequently observed color domains, dividing the dorsal tract in four compartments and the ventral tract in five (Figure 1C). Recording coloration within each herein defined domain showed that despite noticeable trends, each domain could acquire diverse hues and within-feather motifs (Supplementary Table 1), implying that within-feather coloration is acquired independently of color domain position/orientation. Together, this survey shed light on a set of frequent color domains that combined produce most observed color patterns.

**Fig. 1.**
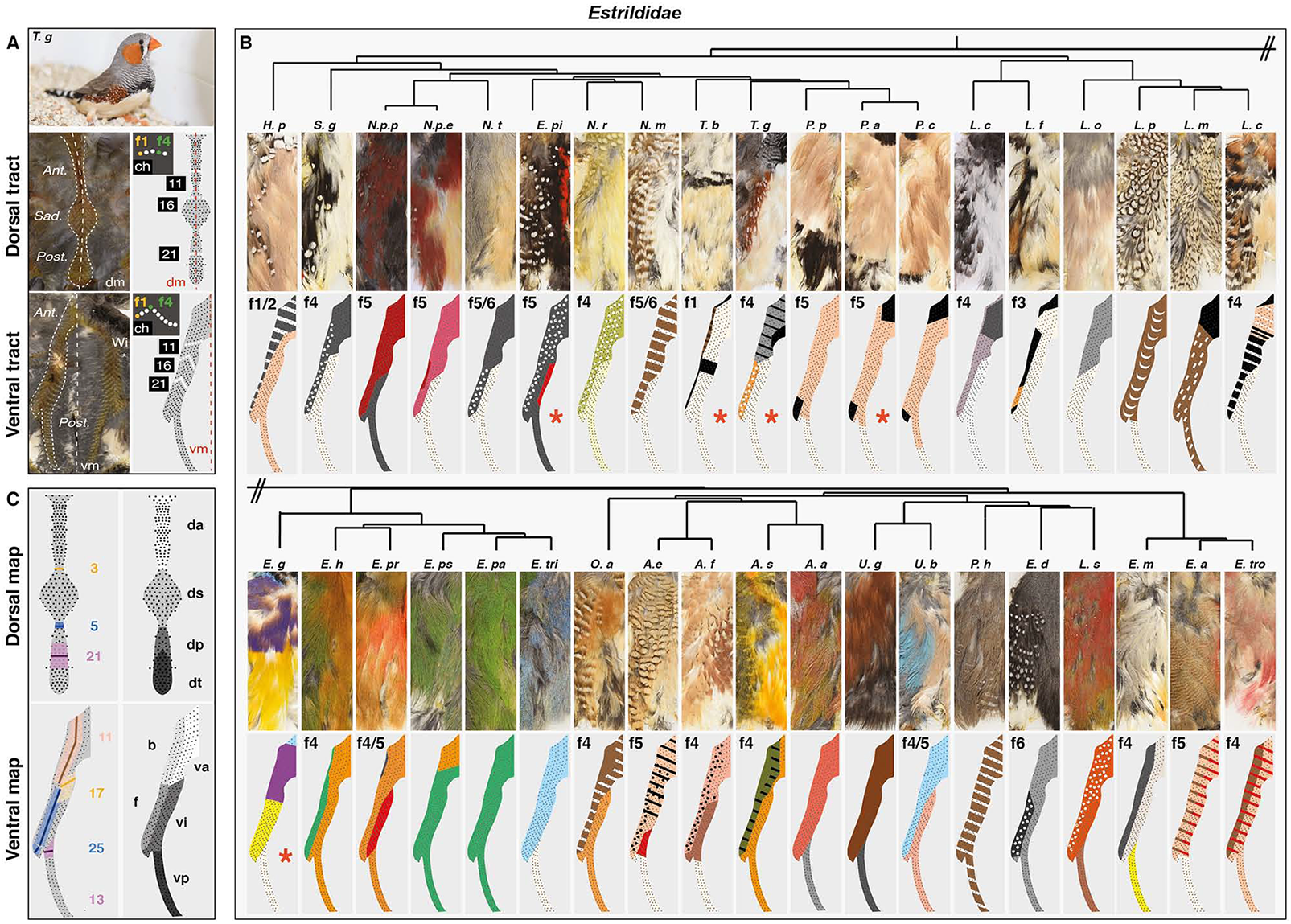
Color domains have recurring position/orientation in *Estrildidae.* **(A)** Flat skin preparations from adult male specimens (here, a zebra finch) allowed visualizing feather implantation in dorsal and ventral tracts (white dotted lines), respectively composed of thin anterior and posterior regions separated by a central “saddle”, and two bilateral sides merging above wing. Schemes show conserved typical tract maps in which feathers (f; black dots) are reproducibly arranged in chevrons (ch11, 16 and 21 are shown) such that they can be assigned a fixed number (f1 and f4 are shown). **(B)** Analyzing coloration on all feathers of ventral tracts (upper panels and see Fig. S2) for 38 *Estrildidae* species representative of the phylogeny (shown schematically and see Fig. S1) produced spatial maps of color domains (bottom panels). If present, the boundary between flank and ventral-most domain is shown with f “X”, indicating on which feather of the chevron it was located. Full species names are listed in Fig S1. **(C)** Comparing the position of color boundaries showed that several are oriented along the tract axis and have recurring position (colored lines, numbers indicate that of species in which the boundary is visible; details of the analysis are shown in Fig. S5). Compiling results allowed creating a reference map with four domains putatively conserved in the dorsum (anterior, saddle, posterior and tail) and five in the ventrum (dorsal-breast, ventral-breast, flank, ventral and posterior). Ant.: anterior, Sad.: saddle, Post.: posterior, dm.: dorsal midline, Wi.: wing, vm.: ventral midline.

To uncover developmental events controlling the formation of frequent domains, we described dynamics of pigment appearance in the zebra finch *Taeniopygia guttata*, in which most frequent color domains are visible, and the owl finch *Taeniopygia bichenovii* and Gouldian finch *Estrildae gouldiae*, which respectively differ in the position of longitudinal and transverse boundaries in the ventrum (see Figure 1). Strikingly in hatchlings of the three species, first visible pigment boundaries were identically located, such that they readily distributed according to the adult pattern in the zebra finch (Figure 2A) but not to the other two species where they were later either shifted or not visible (Supplementary Figure 6). Thus, cryptic boundaries may form even when color domains are apparently lacking in the adult plumage. This strongly suggested frequently observed domains form through common embryonic mechanisms. To seek for those, we described dynamics of tract formation in zebra finch embryos prior to pigment appearance using stains for *β-catenin* transcripts that mark developing feather follicles, or primordia (11). At developmental stage hh30, dorsal primordia first individualized from two *β-cαtenin*-expressing lines extending from neck to tail and fused posteriorly, forming a Y shape. Additional primordia sequentially appeared in a medio-lateral row-by-row wave until tract completion. In the ventral tract, 3-to-4 primordia individualized at hh29 from a droplet-shaped *β-cαtenin*-positive area in the under-wing region, forming a small row that rapidly extends anteriorly and posteriorly, and becomes flanked medially and laterally by additional rows, in a sequential dynamics similar to that observed in the dorsum. At hh37, the same process takes place in the breast and laterally to the leg, and at hh39, all three primordia-forming regions fuse to form the single, continuous surface of the ventral tract (Figure 2B and see (12,13)). In both dorsum and ventrum, timely sequences and overall orientation of primordia emergence did not reflect the position/orientation of color boundaries most-frequently observed in *Estrildidae* (despite some orienting parallel to fusion zones between breast-side and side-leg primordia-forming regions). In addition, dynamics of tract formation were identical in the owl finch in which boundaries differ (Supplementary Figure 7). Thus, mechanisms controlling frequent color domain formation are likely independent of tract emergence. We therefore assessed the lineage of color domains by transplanting early developing zebra finch tissues otherwise known to provide positional information to skin cells into domestic chicken *Gallus gallus* hosts at hh14 (finch eggs being too fragile to sustain grafting). We assessed the effect of grafts in resulting chimeras at hh28 using *β-catenin* for tract spatial reference, and stains for the pigmentation gene *Agouti*. The latter, previously shown to mark presumptive color domains in other species, varies spatially between host and donor species: we observed *Agouti* expression in the dorsal skin of the domestic chicken but not of the zebra finch, and it formed a V-shaped pattern flanking the nascent ventral tract in the domestic chicken, while it covered half of its surface in the zebra finch, marking the position of the future boundary between flank and ventral domains (Figure 2C and Supplementary Figure 8). We grafted trunk-level epithelial somites, which give rise to dorsal tract cells (14) and instruct the position of dorsal *Agouti* expression and colored stripes in poultry birds (Figure 2D, (15)). In the dorsum of hh28 chimeras, *Agouti* expression was absent in the grafted side, similar to control donor zebra finch embryos. Thus, and together with previous results (15), signals from somites are necessary for *Agouti* expression in the dorsum. In the ventrum, *Agouti* displayed identical expression in grafted and un-grafted sides. Thus, somites do not instruct proper spatial pattern of *Agouti* in this region. Similarly, domestic chicken host-like expression of *Agouti* was found on grafted sides both in the dorsum and in the ventrum upon transplantation of trunk-level neural tube halves, showing this tissue does not contribute to *Agouti* pattern establishment. However, when we transferred wing-level portions of lateral plate mesoderm, tissue giving rise to wings (16) and ventral tract cells (17), *Agouti* expression was unchanged in the dorsum of chimeras (which displayed as expected zebra finch donor-like wings), but markedly shifted dorsally in the ventrum with reference to tract position, in a pattern similar to that of donors (Figure 2D and Supplementary Figure 9). Thus, the lateral plate mesoderm instructs the formation of the ventral tract and controls the position of species-specific ventral *Agouti* expression. To assess the effect of the lateral plate mesoderm on ventral pigment distribution in the nascent plumage, we performed long-term grafting experiments using Indonesian Ayam Cemani chicken as hosts: these birds are entirely black due to hyperpigmentation of the skin, feathers, beak, bones, and internal organs (18). Just before hatching at hh46, grafted areas of chimeras displayed zebra finch-like tracts comprised of a darkly pigmented flank and lighter ventral region, separated by a longitudinal boundary whose location was identical to that of control zebra finch individuals (Figure 2E). Thus, the lateral plate mesoderm governs spatially precise formation of color domains in the ventrum. Together, these experiments showed that early landmarks whose lineage gives rise to tracts, namely somites in the dorsum, and the lateral plate mesoderm in the ventrum, also drive –likely through different mechanisms, region-specific coloration.

**Fig. 2.**
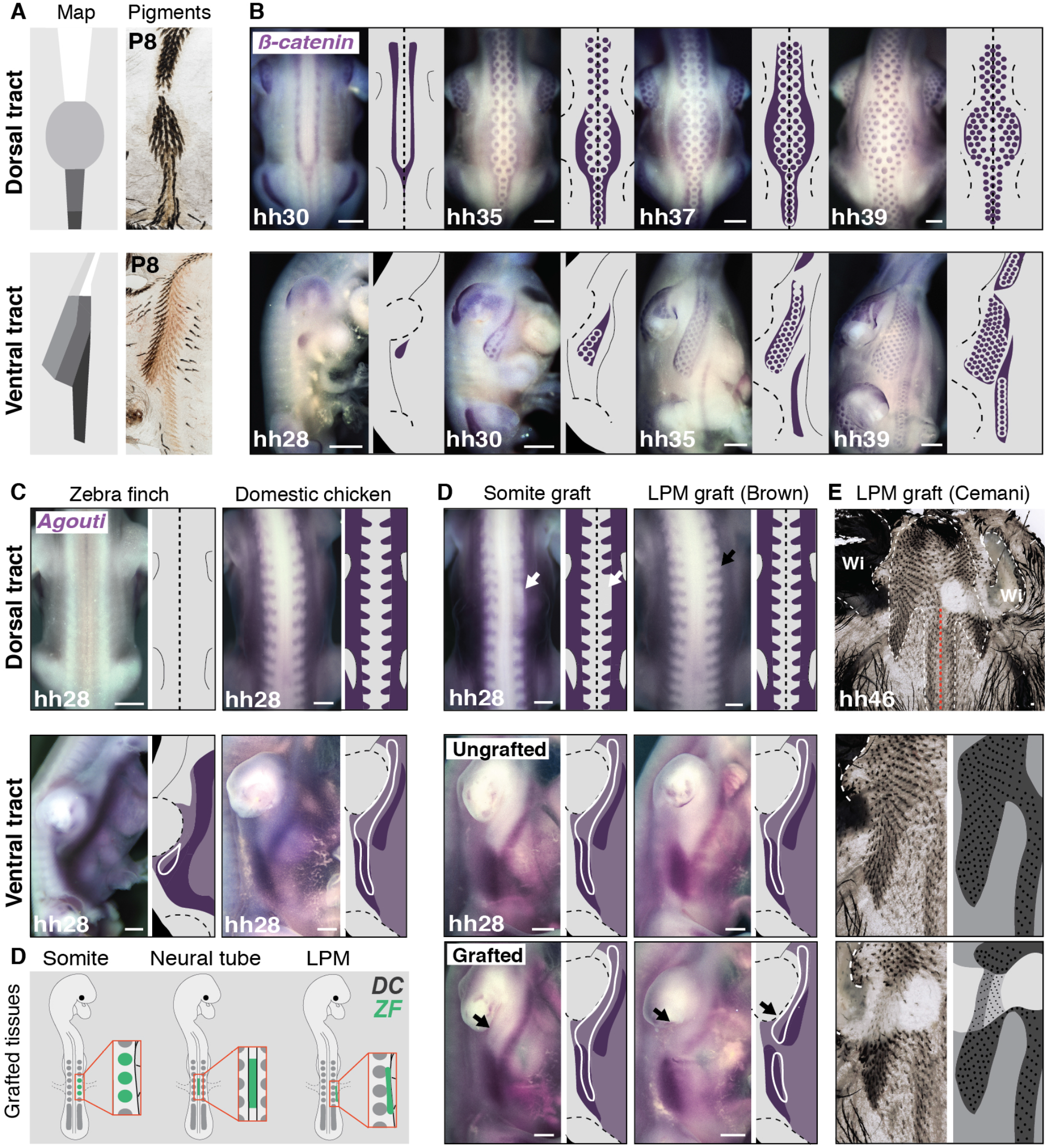
The lateral plate mesoderm instructs the formation of ventral color domains. **(A)** Flat skins preparations in P8 zebra finch individuals showed first visible pigments in the dorsal tract (in two posterior longitudinal domains) and the ventral tract (along the flank-ventral boundary) readily distributed according to the adult pattern (see Figure 1). **(B)** Stains for *β-catenin* transcripts (in purple) and corresponding schematics showed feather primordia emerged in a medio-lateral wave in the dorsal tract of the zebra finch embryo, resulting in the formation of 3 primordia rows in anterior, posterior and tail regions, and 6 in the saddle. Ventral tract primordia emerged in 3 regions progressively extending and fusing to form a continuous pattern. **(C)** Stains for *Agouti* transcripts (in purple) at hh28 showed it was expressed in the dorsum of the domestic chicken but not the zebra finch. In the ventrum, it formed longitudinal bands flanking the nascent tract in the chicken and marking the position of the future flank-ventral boundary in the finch (tract contours are indicated with a white line corresponding to *β-catenin* stains in Fig. S8). **(D)** Schematics representing the transplantation of zebra finch somites, portions of neural tube halves, and of lateral plate mesoderm (in green) in domestic chicken host embryos. **(E)** In somite-grafted Brown strain chimeras at hh28, the expression of *Agouti* (in purple) was absent in the dorsum in the grafted side area, remaining unchanged in the ventrum. In lateral plate mesoderm-grafted Brown strain chimeras, its expression was unchanged in the dorsum but shifted dorsally in the ventral grafted side compared to the control side. Flat skin preparations show that on their grafted side, lateral plate mesoderm-grafted Cemani strain chimeras at hh46 displayed small and pigment-less wing and a zebra finch-like ventral tract comprising a darkly pigmented flank and lighter ventral region separated by a boundary located on f4. DC, domestic chicken; ZF, zebra finch; P: post-embryonic day, LPM: lateral plate mesoderm, Wi: wing. Scale bars: 500 μm.

To identify molecules spatially restricted to color domains, we performed transcriptomics analyses in the zebra finch, whose genome was sequenced and annotated (19). We microdissected embryonic skin regions at hh28 corresponding to the future location of most conserved domains in male embryos, extracted total RNA, and carried through RNA-seq experiments (Figure 3A and see methods). Transcripts levels were analyzed for all combinations of domain pairs. The largest number of differentially expressed genes and highest fold-changes were observed along the dorso-ventral axis. We observed a dorsal enrichment of regulated genes involved in the control of actin cytoskeleton, cell adhesion, matrix-to-receptor interaction, and melanogenesis, expected results as the developing skin is characterized by a dorso-ventral gradient of differentiation –including higher density/activity of pigment cells (12,20). Analyses evidenced high fold-change dorso-ventral differences in transcript levels for several genes belonging to the Wnt and TGF-β signaling pathways, which are involved in skin appendage differentiation (21,22), and homeobox factors, which provide positional identity along body axes (23). Along the antero-posterior axis, fewer genes were differentially regulated, displaying weaker fold-changes, but these were also often identified as pigmentation and cell adhesion genes (Supplementary Figure 10). We retained as candidates genes with at least a 2.5-fold expression change, discarding those up-regulated either in the dorsum or in the ventrum in more than two combinations, as they could reflect the age gradient (e.g. *Zic1, Zic3, Zic4, Foxb1, Crabp1, Hand2).* A high number of homeobox factors met our selection criteria including *Hoxa6, -a9, -b7, -c9, -d3, -d9, Osr2, Pitx2, Shox* and *Tbx18* along the antero-posterior axis, and *Shox, Tbx5, Tbx18, Six2,* and *Irx1* along the dorso-ventral axis. For the latter, some genes were differentially regulated in both dorsal and ventral compartments, such as *Hoxa2, -a3, -a4* that were up regulated anteriorly, *Hoxa7, -a9,* – c*9, Pitx2, Osr2* in intermediate regions, and *Msx2* and *Hoxa11* posteriorly (Figure 3B and Supplementary Figure 11). *Agouti,* which displayed one of the highest fold-changes, was up-regulated in ventral domains, consistent with results described above (see Figure 2). We combined results along both body axes, and found only three genes present in a single putative color domain, namely *Alx4* and *Isl1* in the ventral posterior region and *Tbx18* in the flank. This suggested that color domain formation mostly relies on a combinatory landscape of patterning genes. Together, profiling results provided several candidates potentially marking specific skin regions across both tract axes. We cloned 60 candidate genes (Supplementary Table 2) and qualitatively screened their expression profiles using *in situ* hybridization in the zebra finch embryo at hh28. 37 displayed spatially-restricted expression patterns. In the dorsum, *Hoxa4, Hoxc6, Hoxa7, Ptchd1, Ism1, Pitx2, Alx1, Fzd4* and *Hoxa10* stained staggered skin segments from neck to tail. In the ventrum, *Hoxa4* and *Alx1* encompassed future anterior domains, *Six2*, *Tbx15*, *Tbx18*, and *Pitx2* intermediate regions, and *Irx1* the posterior domain (Figure 3C and Supplementary Figure 12). *Six2* marked the longitudinal flank boundary, and *Tbx15* and *Tbx18* were restricted to the flank domain. In addition, *Osr2* and *Agouti* expressions, detected throughout tract length, marked longitudinal boundaries. Anteriorly, they were restricted to ventral-most regions. Along tract sides, *Osr2* covered the whole nascent tract while *Agouti* marked the flank boundary, its dorsal limit corresponding to the ventral limit of *Tbx15’*s consistent with work in rodents showing that *Agouti* and *Tbx15* interact to control the location of dorso-ventral color boundaries ((24); Supplementary Figure 13). Posteriorly, *Agouti* encompassed the entire future tract while *Osr2* flanked its dorsal border. Remaining candidates were expressed outside of presumptive tracts and/or throughout tract length with varying widths, not spatially correlating with color boundaries (Supplementary Figure 14). Together, quantitative and qualitative expression results showed that while few gene profiles are restricted to single color domains, a limited number are sharp enough in adjacent or overlapping skin portions that they can be combined to delineate future color boundaries, providing a molecular map of presumptive color domains in the zebra finch (Figure 3D).

**Fig. 3.**
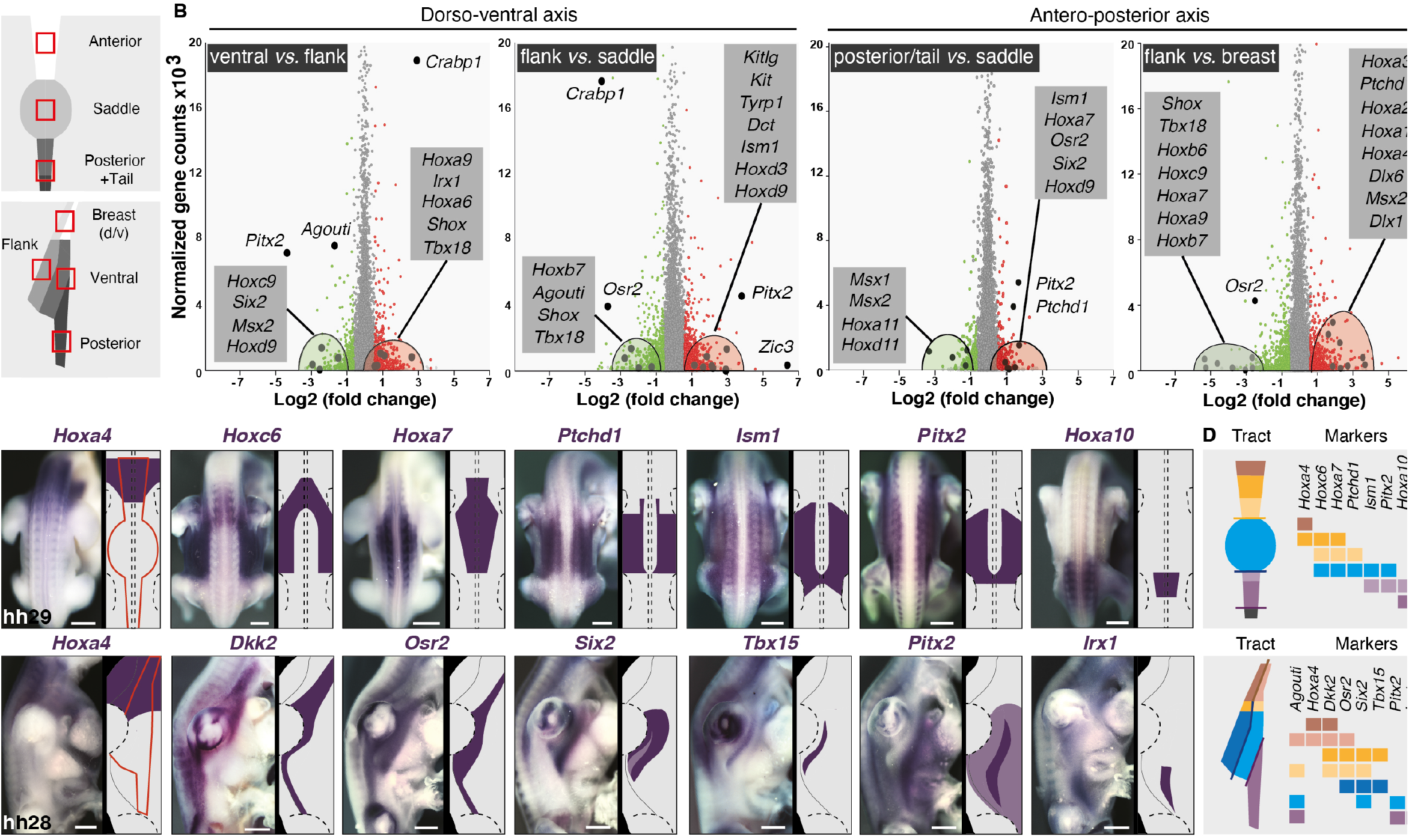
Gene expression profiling identifies markers of putative color domains. **(A)** RNA seq profiling was performed on dissected skin portions corresponding to presumptive color domains (red squares) of zebra finch embryos at hh28. **(B)** Plotting transcript levels as a function of differential expression between pairs of putatively conserved domains along body axes (other combinations are shown in Fig S11) allow identifying significantly up-regulated (in red) and down-regulated (in green) genes. Candidate genes (enlarged dots) are detailed in grey boxes. **(C)** Expression profiles and corresponding schematics are shown for candidate genes indicated in purple in zebra finch embryos upon the emergence of tracts (their putative contours are shown in red). Dotted lines indicate the position of limbs and the neural tube. Other candidates are shown in Fig S14. **(D)** Schematics of compiled markers illustrate that a combination of few genes, together constituting a molecular map, can mark all putatively conserved color domains along body axes in the zebra finch embryonic skin. Scale bars: 500 μm.

To test whether this map is conserved amongst *Estrildidae*, we compared it between species displaying different color boundaries, focusing on the flank/ventral region, where boundaries are most frequent, always longitudinal, and varying in dorso-ventral position. In the zebra finch, the boundary on f4 is marked in the embryo at hh28 by the combination of *Agouti*, *Osr2, Six2, Tbx15,* and *Pitx2* expressions (see Figure 3). In the owl finch, whose flank-ventral boundary is dorsally shifted in f1, *Agouti* and *Tbx15* expressions narrowed down. In the Painted finch *Emblema picta,* in which the ventrum is dark and the flank-ventral boundary separates a periodically ornate from a homogeneous domain, we did not detect *Agouti* and *Tbx15* at that stage. In the long-tailed finch *Poephila acuticauda,* whose ventrum has homogeneous coloration, *Agouti* and *Tbx15* were not complementary, the first having weak, ventrally shifted expression, and the second restricted to a thin band in the dorsal-most part of the flank. Finally, in the Gouldian finch, in which adults have homogeneously yellow flanks but juveniles possess a cryptic boundary identical to that of the zebra finch, *Agouti* and *Tbx15* displayed dorso-ventral extents of expression similar to the zebra finch (Figure 4). Thus, spatially-restricted, complementary expressions of these two markers correspond to species-specific varying positions of color boundaries. Though it did not mark exactly the position of the flank-ventral boundary, *Six2* also varied between species according to differences in the extent of color domains in the side. *Agouti*, *Tbx15* and *Six2* may thus be involved in the control of boundary positioning. By contrast, in all species, *Pitx2* was little changed spatially, and *Osr2* spanned the entire putative flank domain, suggesting they are not involved in the formation of color boundaries. Together, this comparative approach showed that gene expression profiles of the molecular map are largely conserved, with few combined changes rather than modifications in a single gene pattern being sufficient to describe color pattern variation in the flank between chosen species.

**Fig. 4.**
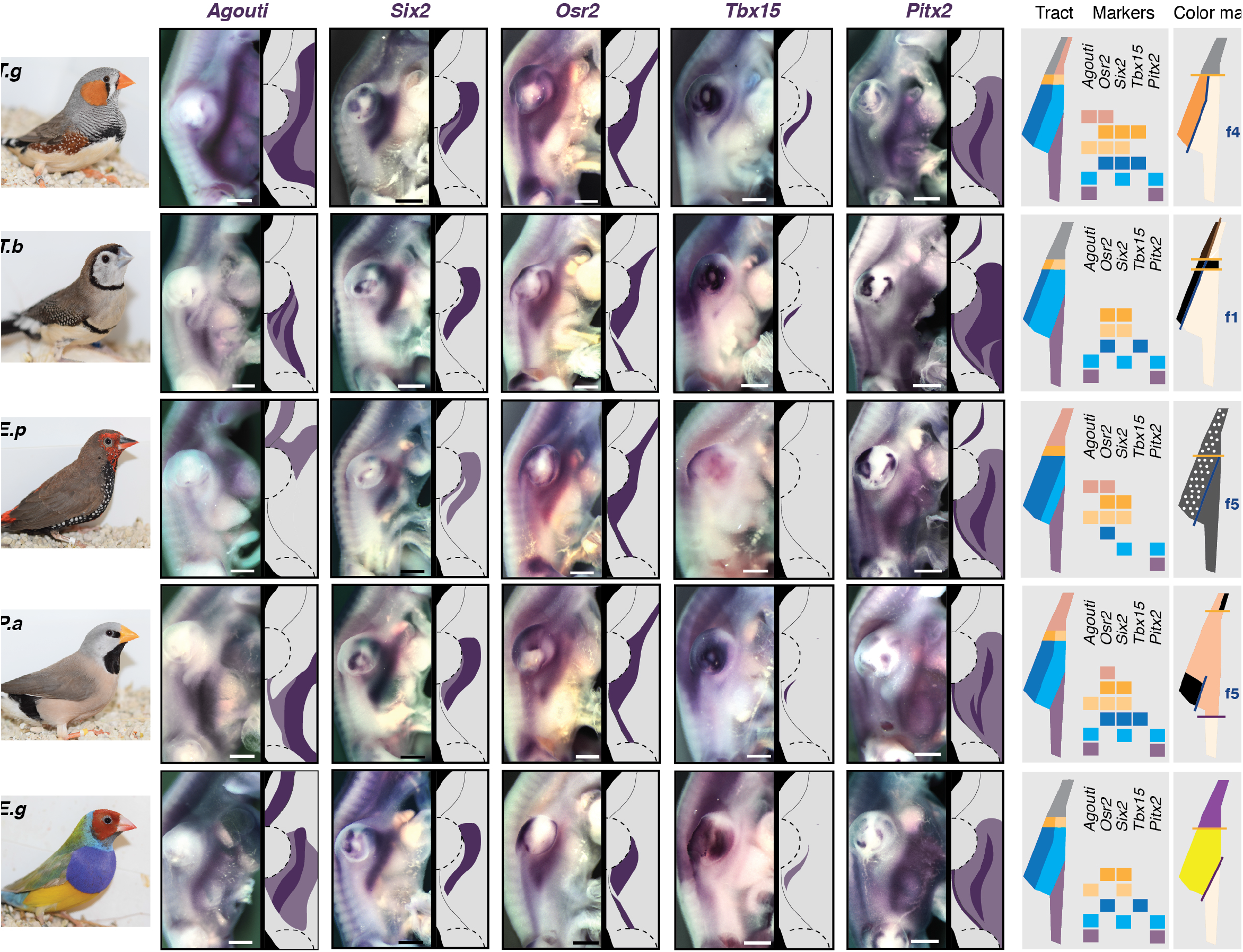
Changes in expression profiles of few markers foreshadow pattern variation”. Expression profiles, corresponding schematics, and marker color codes as defined in Figure 3 are shown for *Agouti*, *Osr2, Six2, Tbx15* and *Pitx2* (in purple) in the ventral embryonic region at hh28 of five species pictured on left panels (species names are listed in Table S1 and below). Compared to the zebra finch, the expression of *Agouti* is shifted dorsally in the owl finch and narrowed-down posteriorly in the Gouldian and the long-tailed finch, corresponding to the position of their flank-ventral boundaries (indicated as f “X” in schematics of adult color maps, right panels). It is absent in the Painted finch, consistent with its dark ventral coloration. The expression of *Tbx15* is also modified according to the adult position of that boundary. By contrast, changes in *Osr2, Six2* and *Pitx2* profiles do not visibly correlate with species-specific color domains. T.g, *Taeniopygia guttata;* T.b, *Taeniopygia bichenovii;* E.p, *Emblema picta;* P.a, *Poephila acuticauda;* E.g, *Erythrura* gouldiae. Scale bars: 500 μm.

Taken together, results of this study evidence common themes underlying apparently extreme variation in color distribution in the adult *Estrildidae* plumage. These spatial trends are due to the establishment of a common “paint box” by early signals from the somite and lateral plate mesoderm. This embryonic template characterized by the spatial combination of genetic markers. A large palette of color pattern variation occurs through limited differential regulation of this combinatory landscape. Extending work to other bird groups and using the molecular map as means to compare and functionally assess candidate pattern-forming factors will be key to decipher whether shared developmental mechanisms shape color pattern diversity in the entire avian phylogeny.

## Acknowledgements

We thank J. Boulton, M. Fiddler, G. Lee and D. Harris for help with bird breeding and adult individual specimen collection, J. Fuchs at the Museum of Natural History of Paris (MNHN) and J. Trimble and Pr. H. Hoekstra at the Harvard University Museum of Natural History for feather specimen collection. We thank Drs. M. Shawkey and J.S. Yeo for help with feather type classification and fruitful exchanges on the project and Drs. B. Prud’homme and N. Haupaix for useful comments on the manuscript.

## Authors contributions

MH performed RNA extraction, cloning of candidate genes, and expression analyses. MH, TLG, NQD, JN and SB collected skin and feather samples and performed phenotypic characterization. RNA-seq data were produced by AM produced and analyzed by Genosplice. MH and CC performed hetero-specific grafting. MH and MM designed experiments and wrote the manuscript.

## Supplementary Materials

### Materials and methods

#### Phenotypic survey of plumage patterns

Flat skin specimens were prepared as previously described (1) from carcasses of adult individuals representing 38 species in the *Estrildidae* family obtained from breeders in Australia (Mike Fiddler), the UK (Dave Harris, John Boulton, Graham Lee) and France (Oisellerie du Temple). Flat skins were imaged using a macro-lens (AF-S Micro NIKKOR 60mm f/2.8G ED) on a D5300 camera (Nikkon). We produced reference maps of dorsal and ventral tracts by plucking out all feathers and recording feather follicle position along longitudinal rows from neck to tail, where contrary to the head region, follicles are arranged in typical chevrons. The dorsal tract is composed of thin anterior and posterior regions separated by a central enlarged “saddle”. The ventral tract is composed of two bilateral sides merging above the wing. In each tract, chevron number is conserved amongst *Estrildidae* species along the antero-posterior axis, and each chevron possesses a conserved number of feathers at specific positions (see Figure 1A; (2)). Color domain maps were obtained by recording at each position feather types classified according to distal hue and motif, all feathers having a proximal grey basis. Hues spanned the entire visible spectrum, created by pigments (i.e, black-brown eumelanin, yellow phaeomelanin, or vivid yellow-to-red carotenoids and porphirins; (3)) or the spatial arrangement of feathers at nano-scale (structural blue-green-purple hues, (4)). In addition, individual feathers display the four periodic motifs described in birds, namely spotted, scaled, barred, or mottled patterns (5).

#### Finch breeding and embryo collection

Male and female individuals of *Taeniopygia guttata, Taeniopygia bichenovii, Erythrura gouldiae, Emblema picta, Peophila acuticauda* were obtained from local suppliers (Oisellerie du Temple, Animotopia) and bred in the bird facility of the Collège de France for collection of fertilized eggs. *Gallus gallus* eggs were obtained from EARL Les Bruyères (Brown strain) and Mes P’tites Cocottes (Cemani strain). Fertilized eggs were incubated at 37°C in Brinsea ova-easy 190 incubators. Embryos were staged according to (6) for finches and (7) for domestic chicken, dissected in PBS, fixed in 4% formaldehyde, dehydrated in EtOH 100% and stored at −20°C.

#### In situ hybridization

*In situ* hybridization experiments were performed as previously described (8). Anti-sense ribo-probes were synthesized from linearized plasmids containing fragments of zebra finch coding sequences for candidate genes (primer sequences are provided in Table S1). The *Tbx5* probe was a kind gift from J. Gros (Pasteur Institute). Double *in situ* hybridizations were performed using ribo-probes tagged with digoxigenin (DIG) or fluorescein RNA-labeling mix (Roche) and sequentially revealed using anti-DIG and anti-Fluorescein alkaline phosphatase antibodies (Roche) followed by reaction on NBT/BCIP (Promega) and Fast Red (Abcam) substrates.

#### Cryosections and Imaging

Embryos were embedded in 15% gelatin and 30% sucrose, sectioned using a CM 3050S cryostat (Leica) and thaw-mounted on Superfrost Plus microscope slides (ThermoFisher Scientific). After washing, sections were mounted in fluoromount (Southern Biotech). Images “ were obtained using a BX53 microscope (Olympus). Whole mount embryos were imaged “ using a MZ FLIII stereomicroscope (Leica) equipped with a DFC 450 camera (Leica).

#### RNA extraction and sequencing

Portions of embryonic brain regions and skin domains corresponding to anterior, saddle, and posterior dorsal regions in the dorsal tract, and to anterior, flank, ventral-most, and posterior regions in the ventral tract (see Figure 3A; n=3 per region for a total of 21 samples), were dissected on ice at stage hh28 (i.e., when the droplet-shaped ventral tract visibly forms in the under-wing region, allowing distinguishing future flank and ventral-most domains). Skin tissues were stored in RNA later solution (Qiagen) at −80°C, brain regions were placed in ATL buffer and used for sex identification through PCR after DNA extraction (Qiagen kit). Sex-specific primers for Z and W chromosomes were designed according to (9): *CHD-W*-For: GGGTTTTGACTGACTAACTGATT; *CHD-W-Rev:* GTTCAAAGCTACATGAATAAACA; *CHD-Z*-For: GTGTAGTCCGCTGCTTTTGG; *CHD-Z*-Rev: GTTCGTGGTCTTCCACGTTT. Total RNA was extracted from male skin tissues (Qiagen kit) and if of sufficient quality (Bioanalyzer: RIN > 7), used to generate cDNA and RNA sequencing libraries at the genomic core facility of the Ecole Normale Supérieure (Paris). In short, 10 ng of total RNA were amplified and converted to cDNA using SMART-Seq v4 Ultra Low Input RNA kit (Clontech). An average of 150 pg of amplified cDNA was used for library preparation following Nextera XT DNA kit (Illumina). Libraries were multiplexed by 21 on 1 high-output flow cells. A 75 bp paired-end read sequencing was performed on a NextSeq 500 device (Illumina). A mean of 19 ± 2 million reads passing Illumina quality filters was obtained for all 21 samples.

#### RNA-seq data analysis

Sequencing, data quality, reads repartition (e.g., for potential ribosomal contamination), and inner distance estimation were performed using FastQC, Picard-Tools, Samtools and rseqc. Reads were mapped using STARv2.4.0 (10) on the taeGut1 Zebra finch genome assembly. Gene expression regulation study was performed as previously described (11). Briefly, for each gene present in the Zebra finch FAST DB v2017_1 annotations, reads aligning on constitutive regions (that are not prone to alternative splicing) were counted. Based on these read counts, normalization and differential gene expression were performed using DESeq2 (12) on R (v.3.2.5). Only genes expressed in at least one of the two compared experimental conditions were further analyzed. Genes were considered as expressed if their rpkm value was greater than 92% of the background rpkm value based on intergenic regions. Results were considered statistically significant for uncorrected p-values ≤ 0.05 and fold-changes ≥ 1.5. Among 18 207 annotated genes, 2999 were regulated (i.e., fold-change ≥ 1.5 with p-value ≤ 0.05). 2150 were uniquely up-regulated genes, 2080 were uniquely down-regulated genes. Clustering and heatmaps have been performed using “dist” and “hclust” functions in R, Euclidean distance and Ward agglomeration methods. GenoSplice performed all RNA-Seq data analyses (www.genosplice.com).

#### Pathway/Gene Ontology (GO) analysis and Transcription Factor analysis

Analyses for enriched GO terms, KEGG pathways and REACTOME pathways were performed using DAVID Functional annotation Tool (v6.8). GO terms and pathways were considered as enriched if fold enrichment was ≥ 2.0, uncorrected p-value ≤ 0.05 and minimum number of regulated genes in pathway/term ≥ 2.0. Results were obtained by compiling analyses performed using either all regulated genes or only up-regulated genes or down-regulated genes.

#### Hetero-specifìc grafting

Grafting experiments were adapted from procedures described in (1). For zebra finch donor tissue preparation, we dissected adjacent somites 18 to 21, portions of neural tube halves (at the level of somites 18 to 21) or of lateral plate mesoderm (at the level of somites 16 to 20) at stage hh14. Donor tissues were transplanted in hh14 domestic chicken hosts in which equivalent tissues had been previously removed. Embryos were visualized *in ovo* by injecting Indian ink (Pelikan) in PBS underneath embryos. Chimeras were incubated for 4 or 18 days and prepared for *in situ* hybridization or flat skins as described above.

**Table S1.**
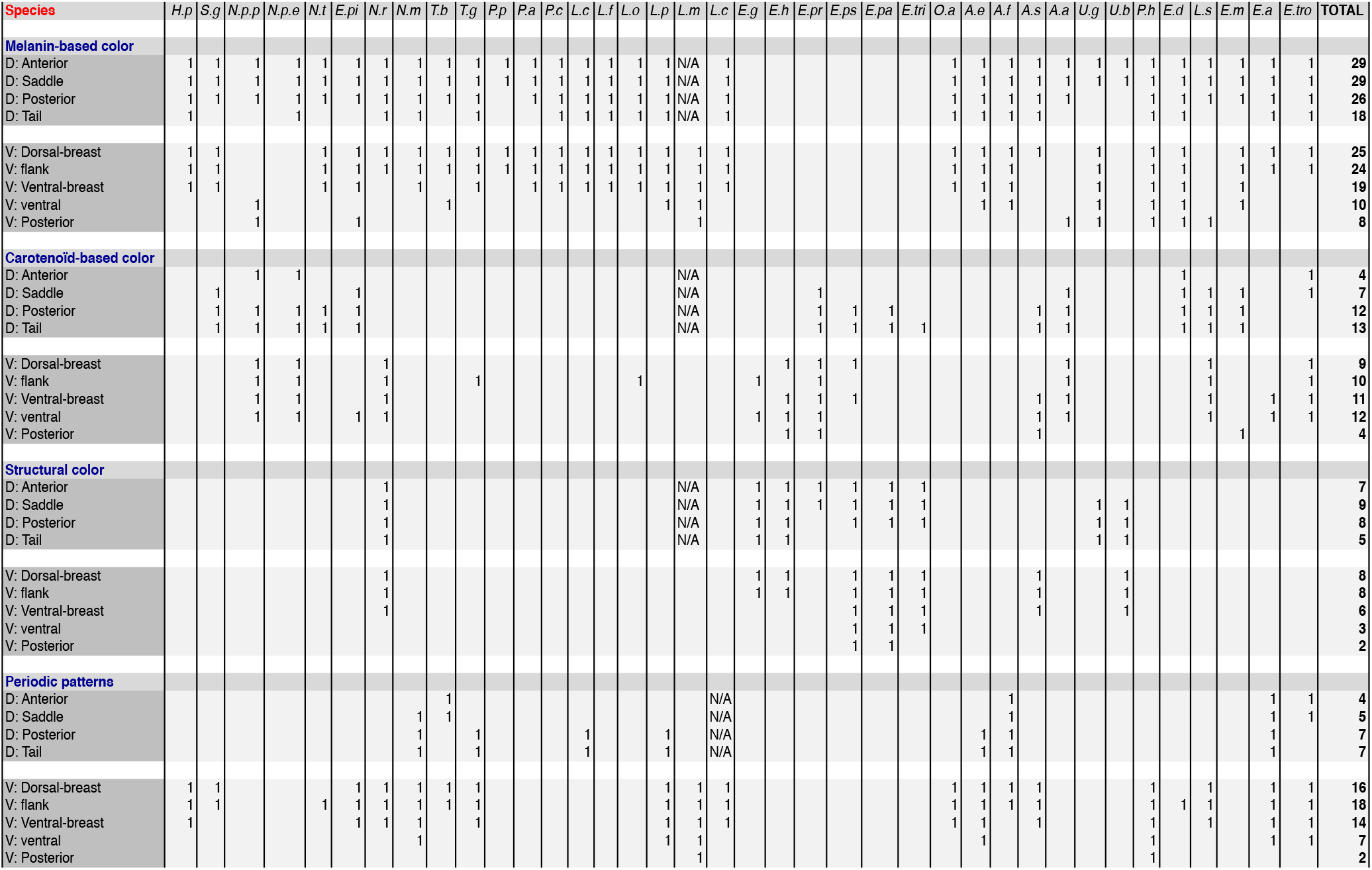
Feather types in conserved color domains.

Recording the presence (noted with 1) of melanin-based, carotenoïd-based, structural, or periodic coloration on individual feathers implanted in frequent color domains (anterior, saddle, posterior and tail regions in the dorsum; dorsal-breast, ventral-breast, flank, ventral, and posterior regions in the ventrum) showed that despite trends, domains display diverse hues and motifs. Ventral domains frequently displayed periodic motifs (20 species, mostly in anterior regions) and light melanin (27 species) or carotenoïd-based (12 species) coloration, but some possessed dark (6 species) or structural (8 species, mostly anteriorly) colors. Dorsal domains most often displayed brown-grey coloration (31 species), but we also observed carotenoid-based hues (14 species, mostly in posterior regions), structural colors (9 species, mostly anteriorly), and periodic motifs (7 species). Species names abbreviated as in Fig. S1.

**Table S2.**
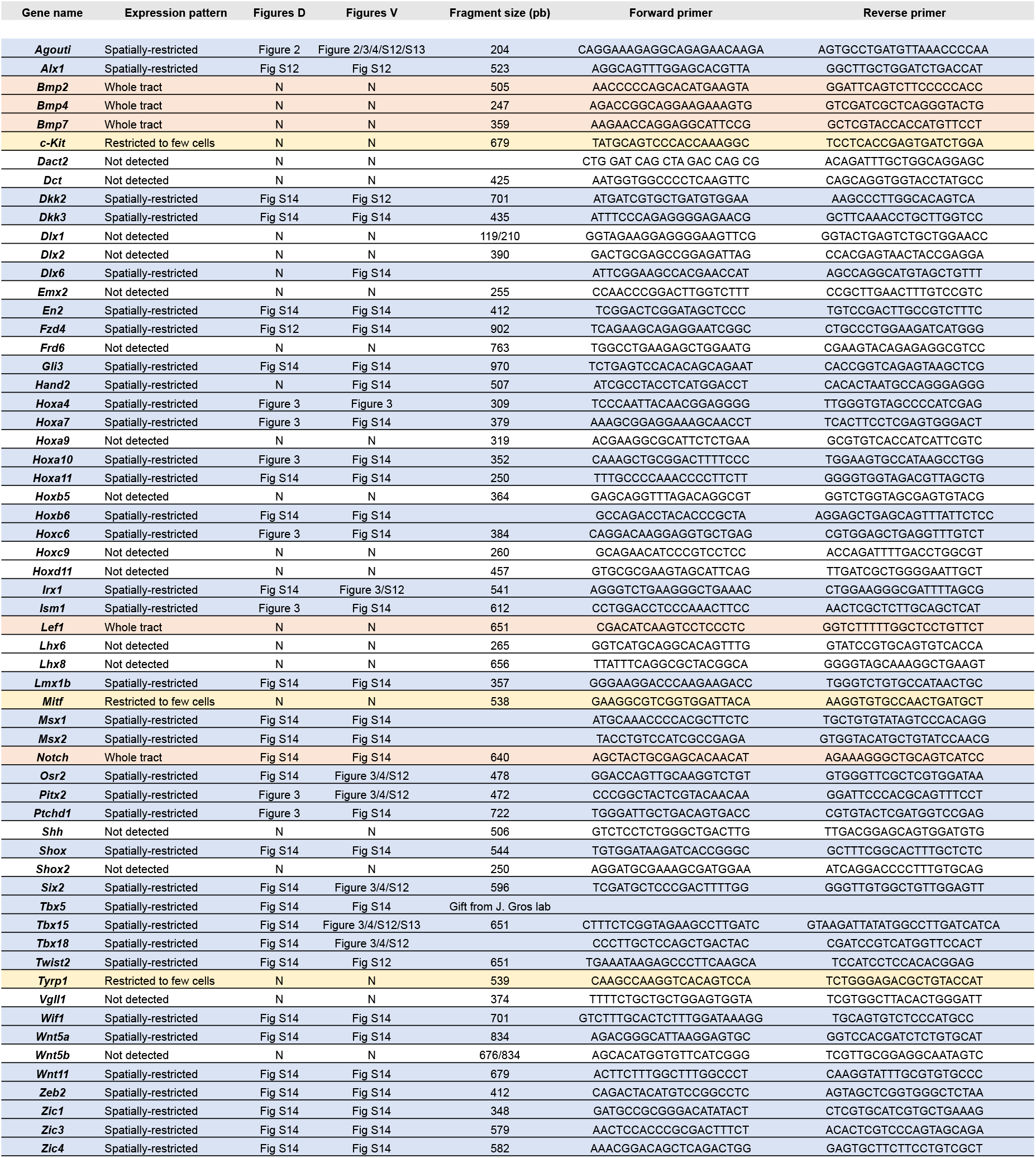
Expression profiles and primers for studied candidates.

We qualitatively assessed the expression of 60 candidate genes (left column) extracted from RNAseq profiling. We cloned anti-sense probes for 59 candidates using primer sequences listed in right columns; the *Tbx5* probe was a kind gift from J. Gros laboratory. For each gene we indicate whether it was spatially-restricted within tracts (in blue), found only in a few cells (in yellow), present throughout tracts (in pink), or not detected (in white), and the manuscript Figures where dorsal (D) and ventral (V) expression data is presented.

**Fig. S1.**
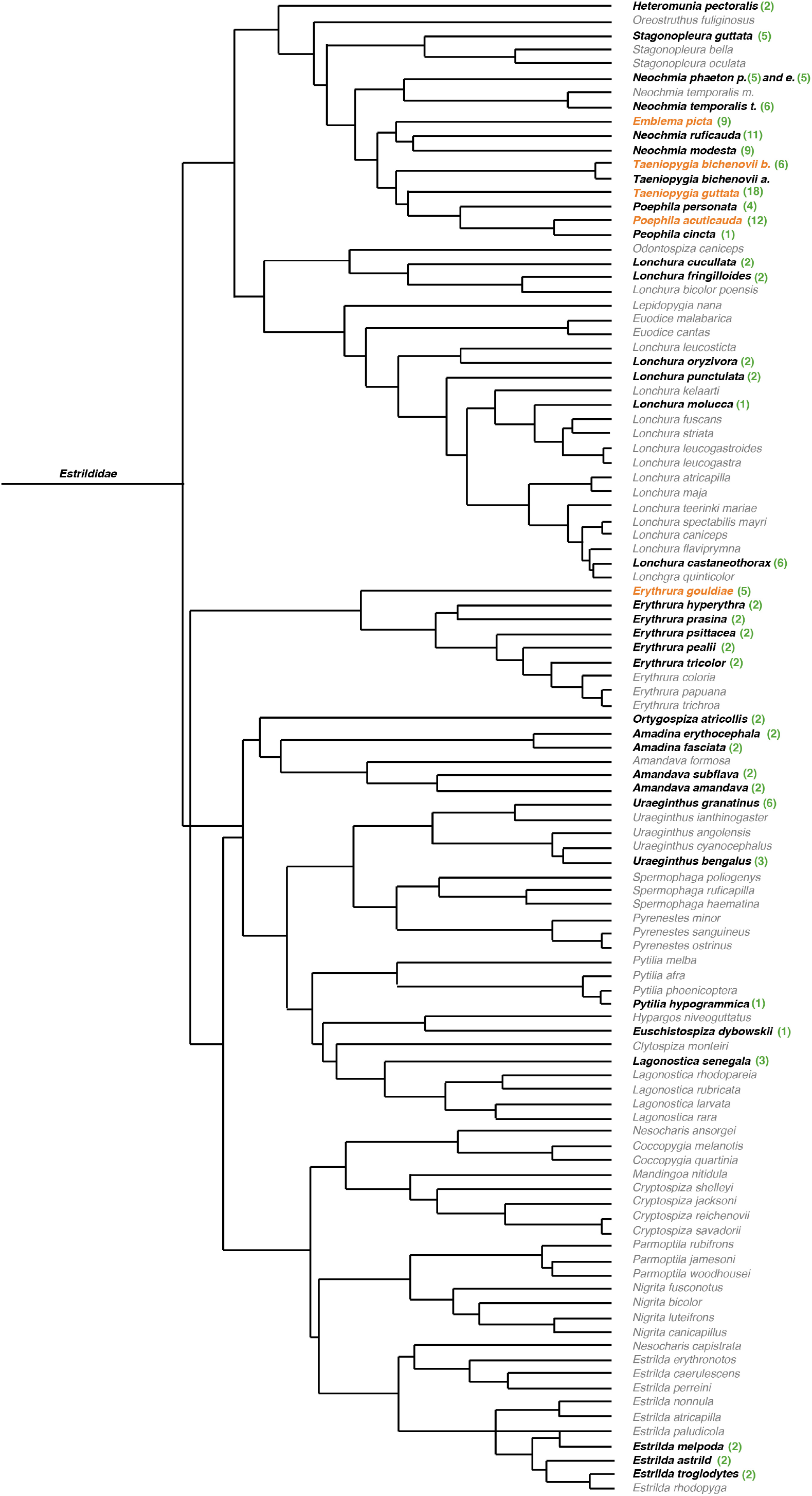
Phylogeny of the *Estrildidae* family. A 105-species phylogeny of the *Estrildidae* family modified from (13) highlights the position of the 38 studied-species (in bold) shown in Figure 1 and Fig S3, namely *Heteromunia pectoralis* H.p, *Stagonopleura guttata* S.g, *Neochmia phaeton phaeton* N.p.p, *Neochmia phaeton evangelinae* N.p.e, *Neochmia temporalis* N.t, *Emblema picta* E.pi, *Neochmia ruficauda* N.r, *Neochmia modesta* N.m, *Taeniopygia bichenovii bichenovii* T.b, *Taeniopygia guttata* T.g, *Poephila personata* P.p, *Poephila acuticauda* P.a, *Poephila cincta* P.c, *Lonchura cucullata* L.c, *Lonchura fringilloides* L.f, *Lonchura oryzivora* L.o, *Lonchura punctulata* L.p, *Lonchura molucca* L.m, *Lonchura castaneothorax* L.c, *Erythrura gouldiae* E.g, *Erythrura hyperythra* E.h, *Erythrura prasina* E.pr, *Erythrura psittacea* E.ps, *Erythrura paeli* E.pa, *Erythrura tricolor* E.tri, *Ortygospiza atricollis* O.a, *Amandina erythrocephala* A.e, *Amandina fasciata* A.f, *Amandava subflava* A.s, *Amandava amandava* A.a, *Uraeginthus granatina* U.g, E.d, *Uraeginthus bengalus* U.b, *Pytilia hypogrammica* P.h, *Euschistospiza dybowskii* E.d, *Lagonostica senegala* L.s, *Estrilda melpoda* E.m, *Estrilda astrild* E.a, *Estrilda troglodytes* E.tro. Numbers of analyzed samples per species are shown in green, species chosen for developmental analyses are shown in orange.

**Fig. S2.**
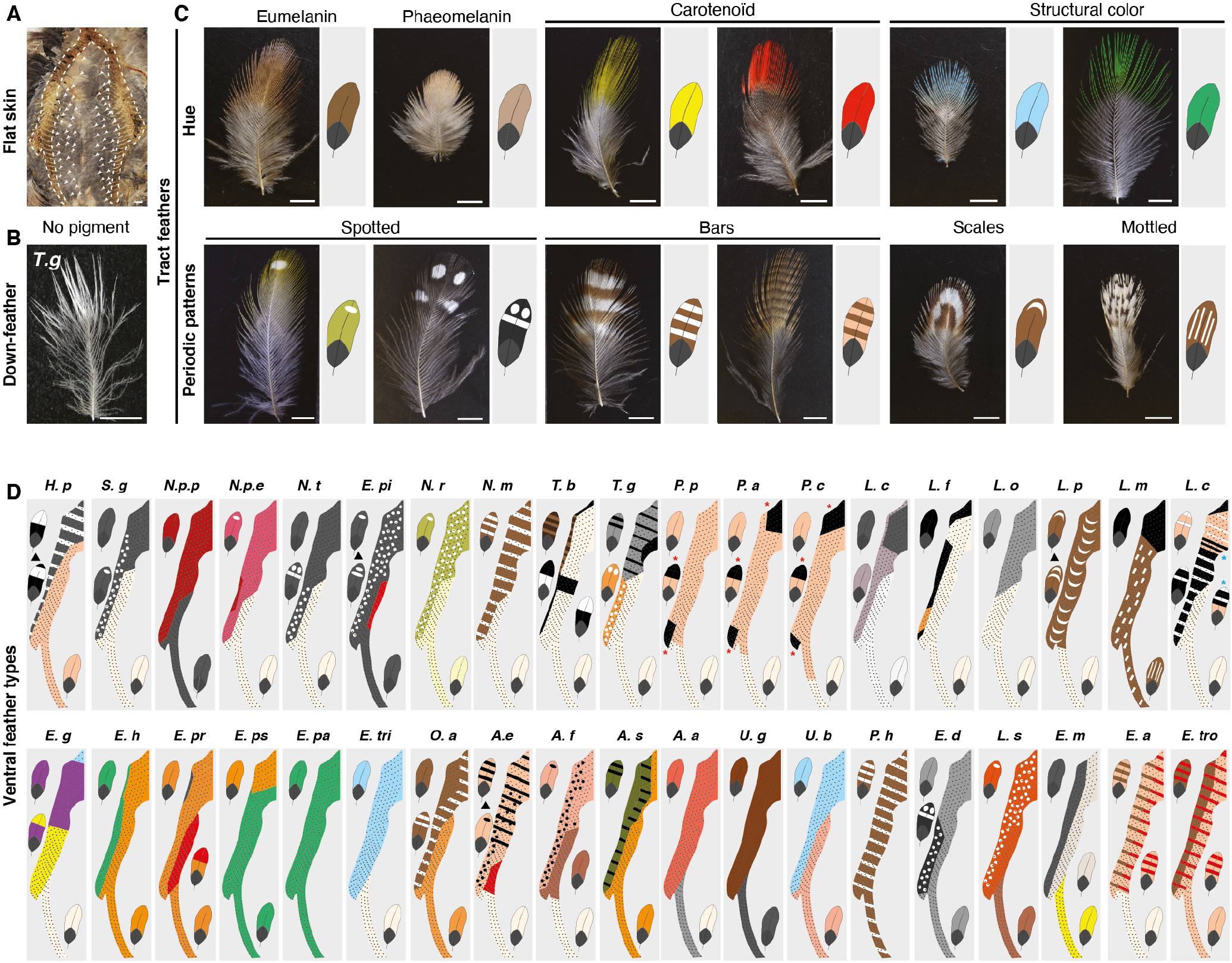
Feather type classification in the *Estrildidae* family. **(A)** Ventral flat skin preparation of an adult male zebra finch showing sparse spatially random distribution of feathers (arrowheads) outside of tracts (dotted lines). Scale bar: 2 mm. **(B)** Feathers outside of tracts have downy structure and white distal coloration. **(C)** Tract feathers and corresponding schematics exemplifying classification according to distal coloration produced by eumelanin (i.e. brown-to-black), phaeomelanin (yellowish-orange), carotenoids (vivid yellow-to-red), structure (blue, green or purple), and/or melanin-based periodic motifs, namely spots, scales, bars and mottled patterns (5). All feathers have a proximal grey basis. Scale bars: 0,3 cm. **(D)** Feather coloration is represented schematically for observed ventral domains in all studied species, providing color-codes on tract maps also shown in Figure 1. Species names are abbreviated as detailed in Fig. S1.

**Fig. S3.**
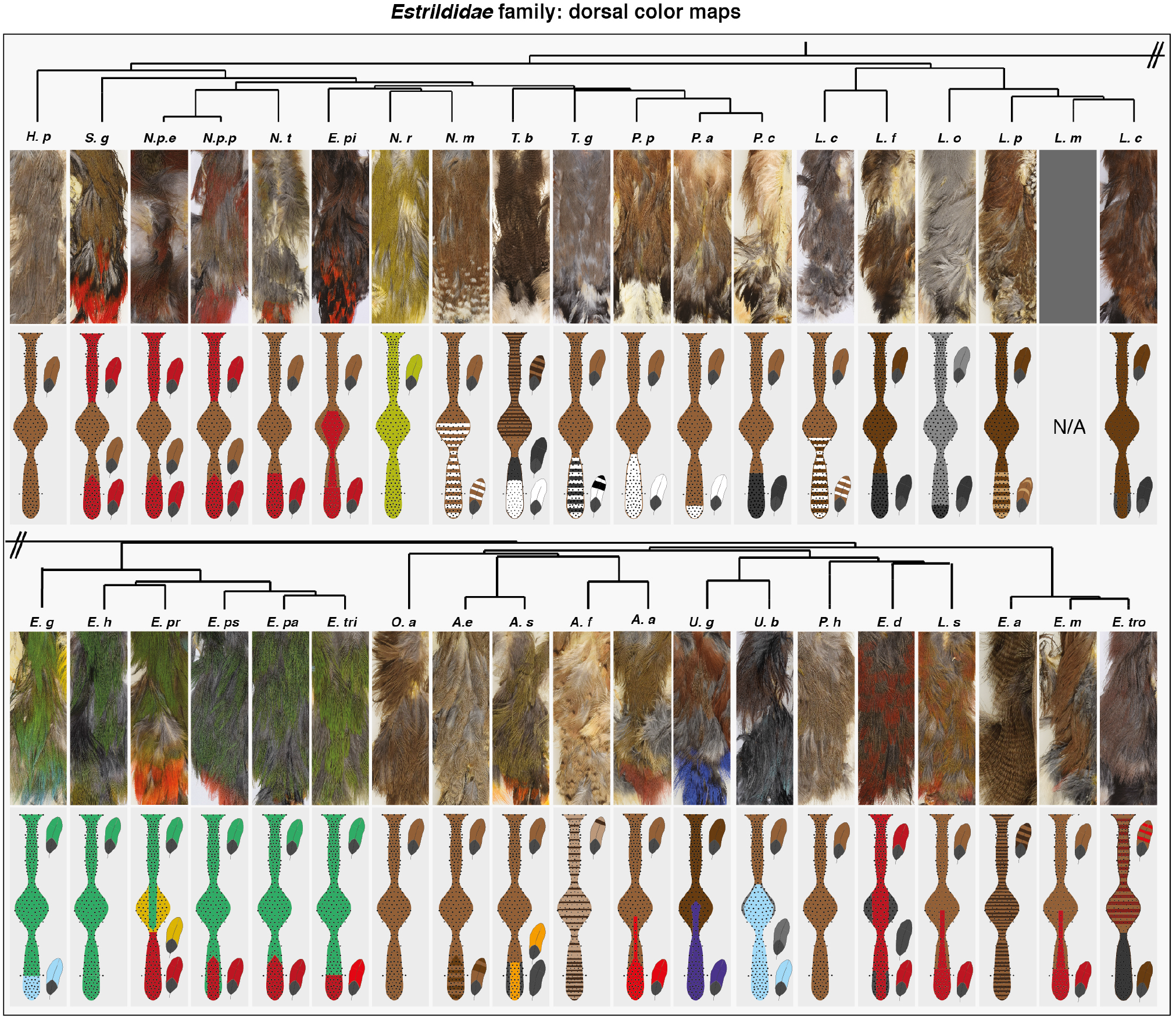
Dorsal color domains in the *Estrildidae* family. Feather coloration recorded on flat skins (upper panels) and color-coded according to methods described in Fig S2 is represented schematically (bottom panels) for observed dorsal domains in all studied species (but for L.m for which flat skin specimens were not available). Species names are abbreviated as detailed in Fig. S1.

**Fig. S4.**
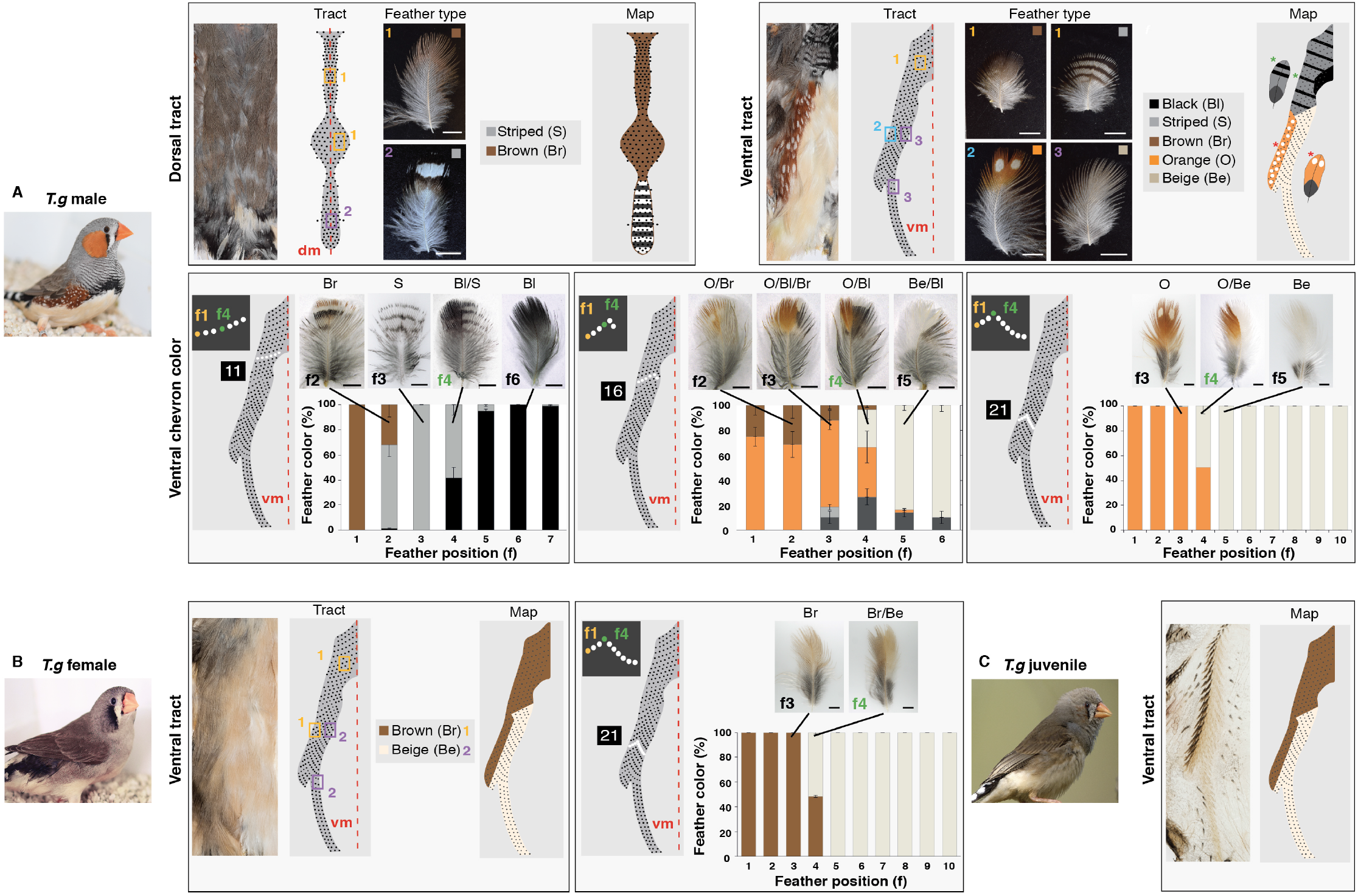
Color pattern characterization in the zebra finch. **(A)** In zebra finch males, recording feather types at each position of dorsal and ventral flat skin preparations (corresponding maps are represented schematically) allow building a typical color domain map (upper panels). Quantification methods are specified for the ventrum (bottom panels): percentages of black, brown, orange, beige and periodically-colored zebra finch ventral tract feathers along chevrons 11 (breast level), 16 (wing level), and 21 (trunk level) show that while color domains may be separated by gradients of pigment type or periodic motifs in the wing and chest regions, overall mean boundary positions are highly reproducible between individuals (n=5). The longitudinal boundary separating the flank from the ventral domain, most often observed in *Estrildidae* (see Figure 1 and S3), is also the sharpest: in the zebra finch it precisely locates on f4 throughout tract length, such that this feather displays a split pattern (i.e., dorsal half orange and ventral half beige). **(B)** In zebra finch females, the flank is brown instead of orange, and f4 displays a brown dorsal half and beige ventral half (n=3). **(C)** In zebra finch juveniles, who display a grey flank, color boundaries have the same position than in adult individuals. Error bars: SEM. dm: dorsal midline, vm: ventral midline. Scale bars: 2 mm.

**Fig. S5.**
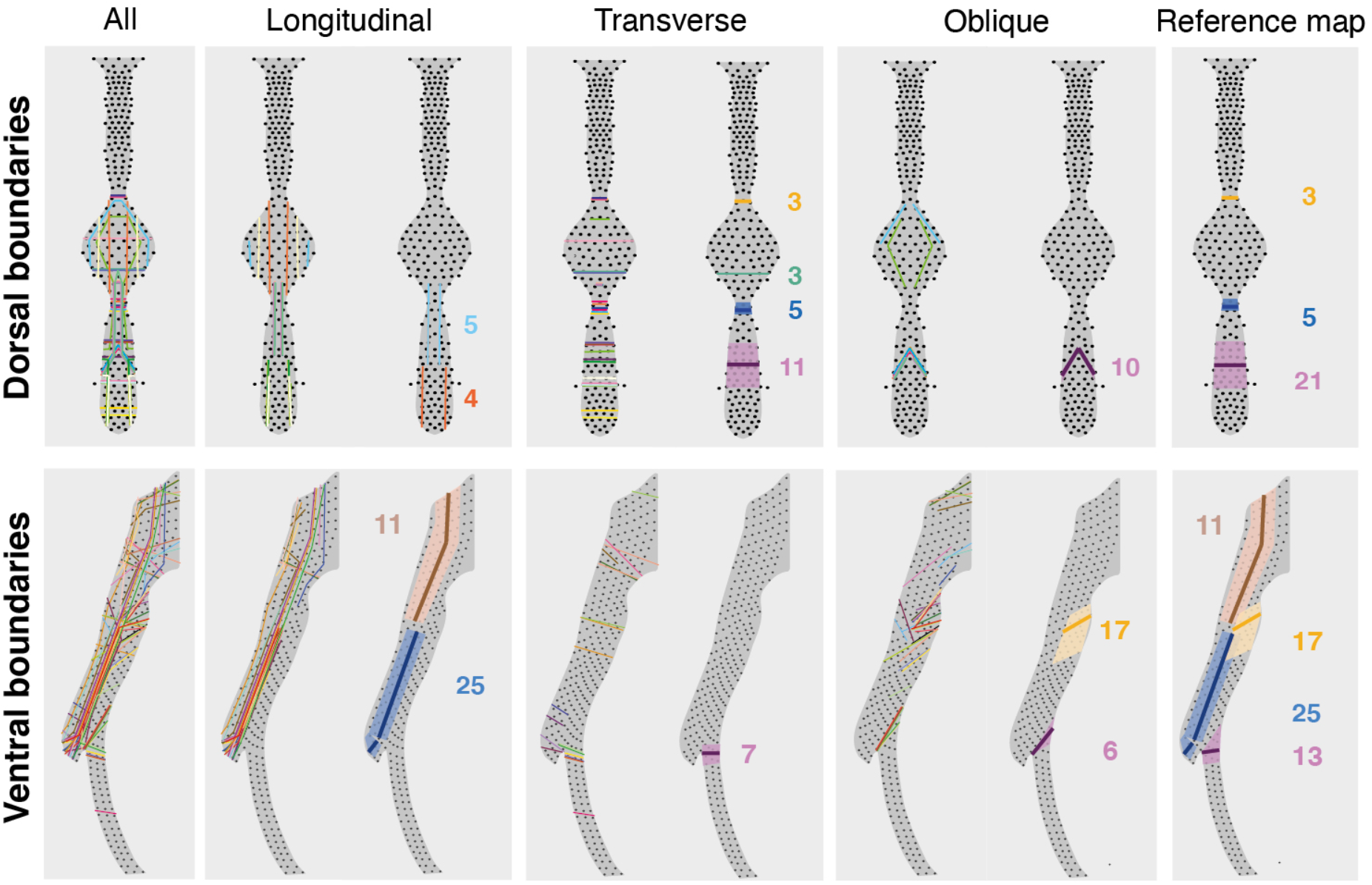
Compilation of color boundaries in the *Estrildidae* family. Observed boundaries with longitudinal, transverse, and oblique orientation with regards to tract axis are represented on dorsal and ventral tract maps. Respectively 76% and 74% of dorsal and ventral boundaries were parallel to tract axes (i.e., not oblique). Longitudinal boundaries were sharp while transverse or oblique boundaries frequently formed gradients. Boundaries were not continuous between dorsal and ventral tracts. Right panels show boundaries observed most frequently (colored numbers represent that of species displaying boundaries located within the corresponding area, lines represent mean boundary position).

**Fig. S6.**
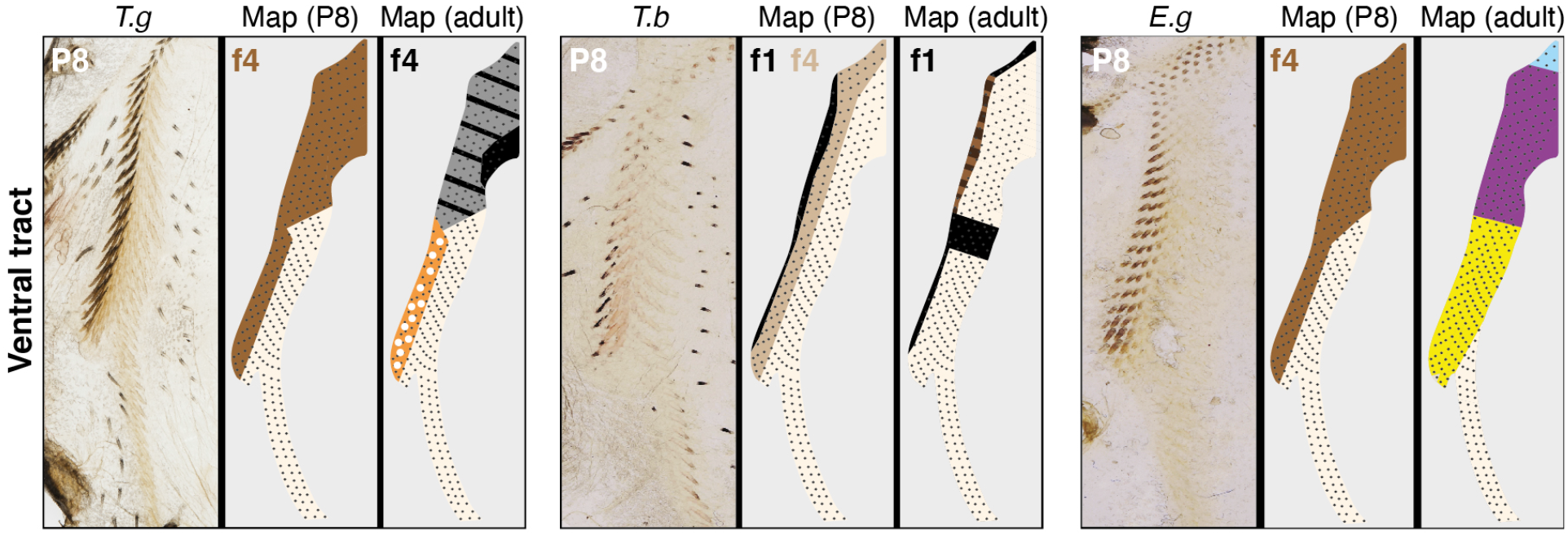
Juvenile color boundaries in the zebra, owl and Gouldian finches. Comparing flat skins preparations and corresponding schematics of ventral skin regions at P8 with color maps of adult individuals of zebra finch *(T.g),* owl finch *(T.b)* and Gouldian finch *(E.g)* shows that while boundaries between flank and ventral domains located on f4 in all juveniles, their position differs in adults (i.e., on f4 in *T.g,* on f1 in *T.b* and absent in *E.g*). P, post-embryonic day.

**Fig. S7.**
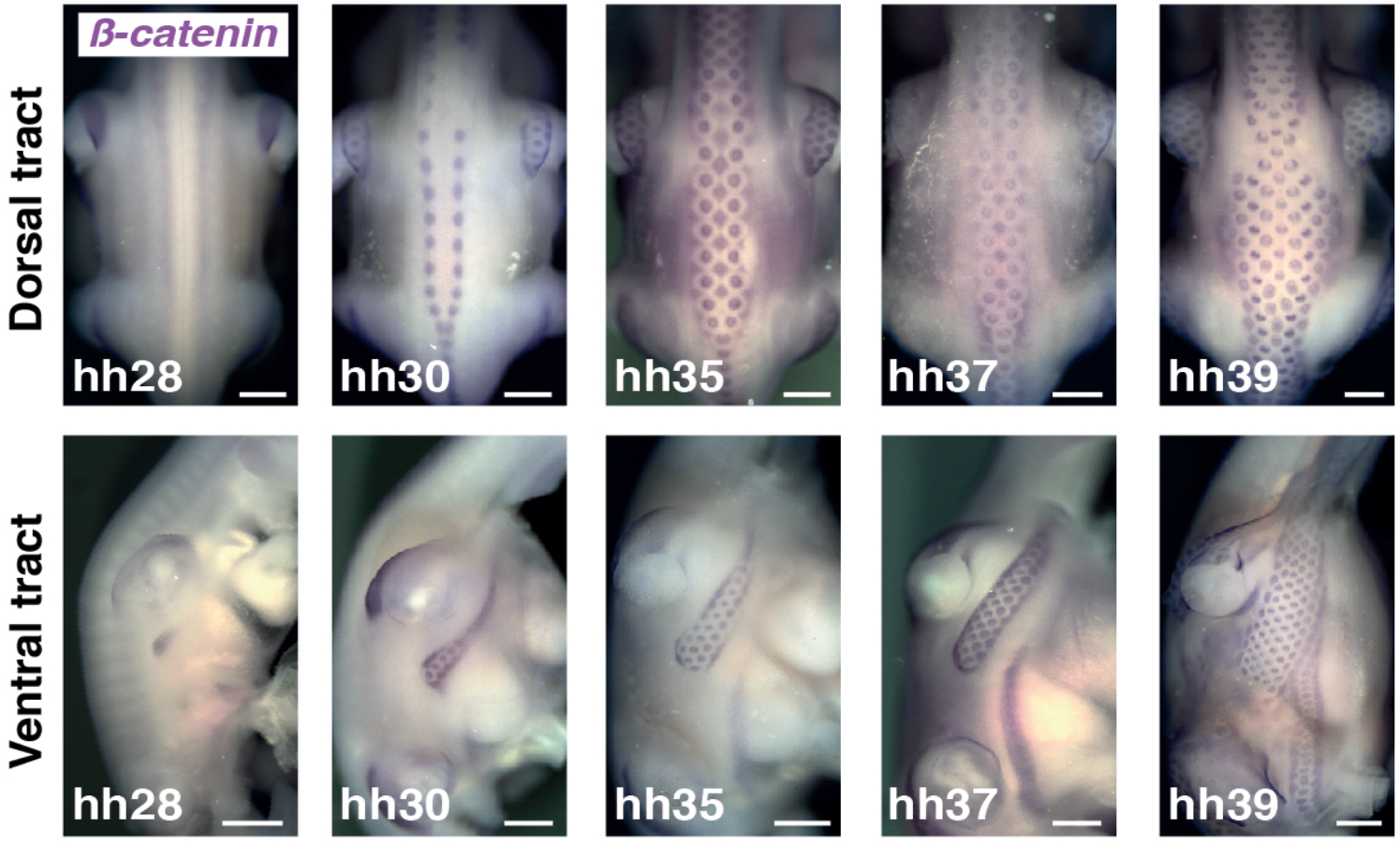
Tract formation dynamics in the owl finch. Stains for *β-catenin* transcripts mark developing feather primordia during ventral and dorsal tract formation in the owl finch *Taeniopygia bichenovii.* The timely sequence of primordia emergence is identical to that described in zebra finch embryos, consistent with previous work showing tracts are conserved throughout the *Estrildidae* family (2). Scale bars: 500 μm.

**Fig. S8.**
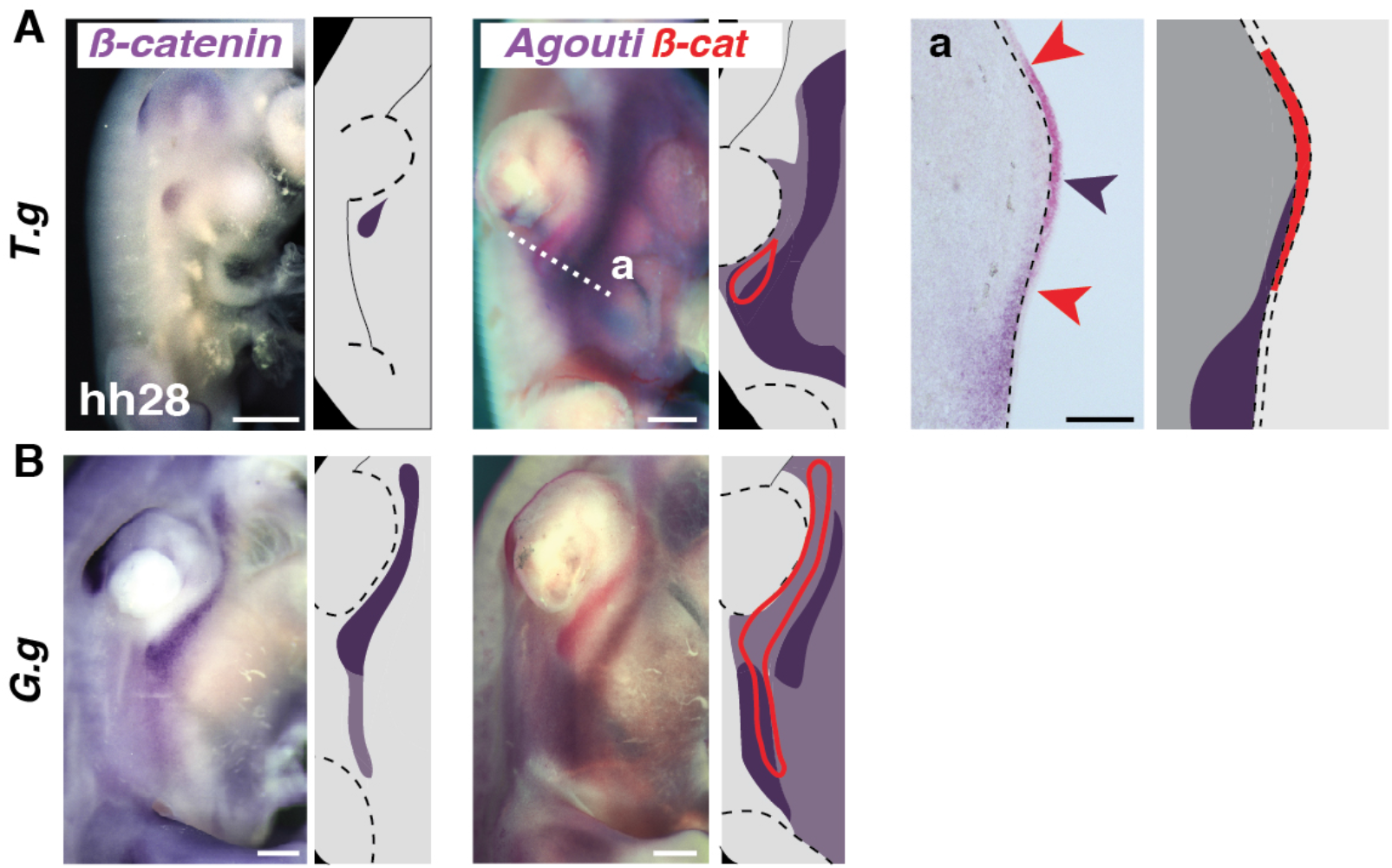
Extent of *Agouti* expression in zebra finch and domestic chicken embryos. **(A)** Double *in situ* stains in zebra finch embryos at hh28 show that *Agouti* (in purple) forms a longitudinal band overlapping with the forming tract marked by *β-catenin* (in red, and red line in schematics). On transverse sections (a; white dotted line), the dorsal limit of *Agouti* expression (purple arrow in schematics) marks the position of the future boundary between flank and ventral domains (stained with *β-catenin*, red arrows). *Agouti* is present in the dermis while *β-catenin* stains the epidermis (the limit between skin layers is shown with black dotted lines). **(B)** In domestic chicken embryos at hh28, *Agouti* does not overlap with *β-catenin* in the flank-ventral region. T.g, *Taeniopygia guttata;* G.g, *Gallus gallus.* Scale bars: 500 μm (section: 100 μm).

**Fig. S9.**
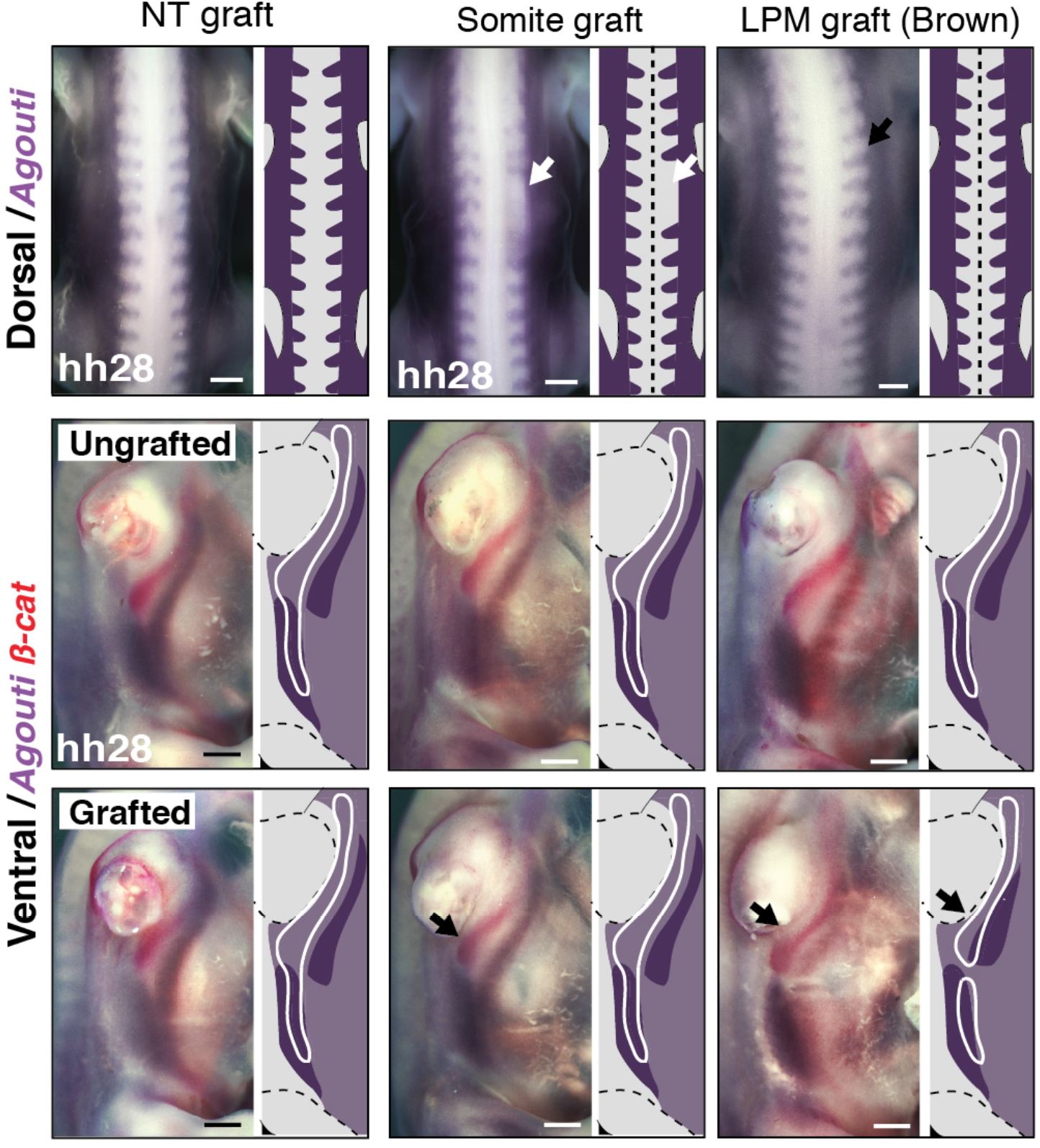
Neural tube grafts and double stains. In neural tube-grafted Brown strain chimeras at hh28, chicken-like expression of *Agouti* (in purple) was maintained in both dorsum and ventrum in the grafted side area. For all grafted chimeras (neural tube, somite and LPM), *β-catenin* (in red, and white lines in corresponding schematics) marks the developing ventral tract. NT, neural tube; LPM, lateral plate mesoderm. Scale bars: 500 μm.

**Fig. S10.**
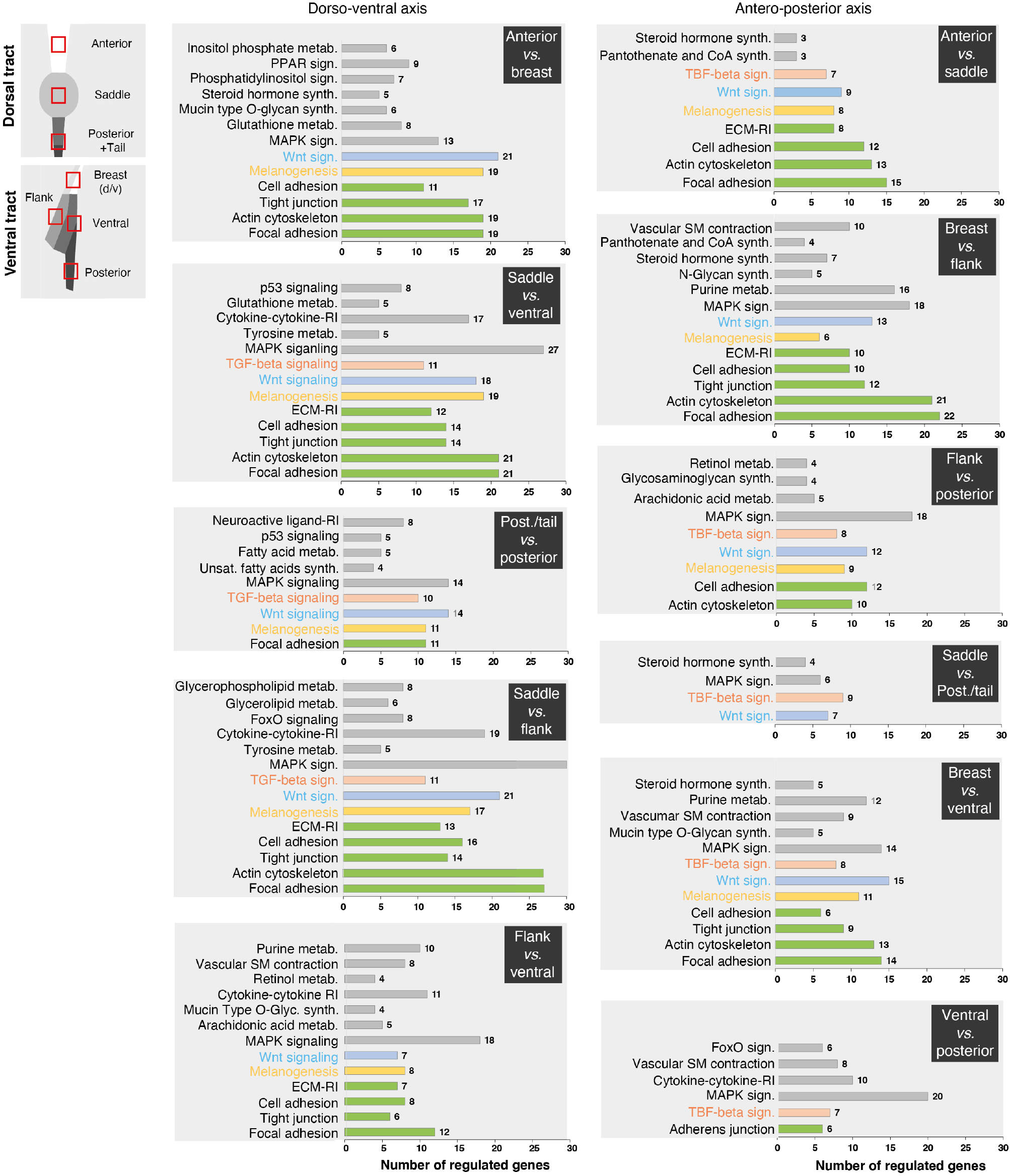
KEGG pathways analyses for all domain pairs. KEGG pathways results displayed for all combinations of domain pairs along dorso-ventral and antero-posterior axes show an enrichment of regulated genes involved in tissue architecture (in green), melanogenesis (in yellow), the Wnt signaling pathway (in blue), the TGF-β signaling pathway (in orange), and other KEGG terms (in grey). Post., posterior; metab., metabolism; sign., signaling; synth., synthesis; ECM, extra-cellular matrix; RI, receptor interaction; unsat., unsaturated.

**Fig. S11.**
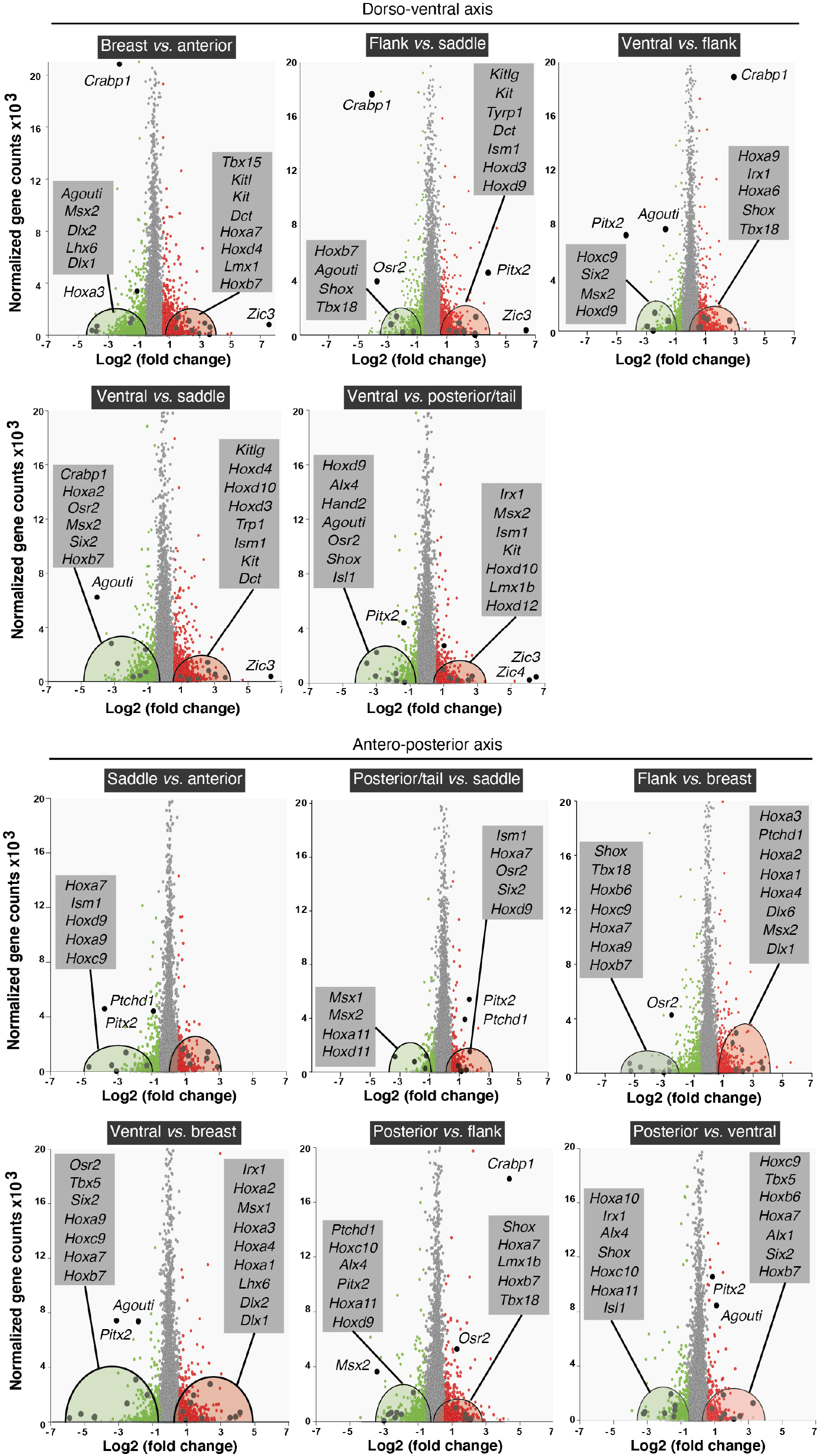
RNA-Seq transcript levels. Normalized genes counts were plotted as a function of differential expression (log2 foldchange) for combinations of domains in pairs. Genes showing significant differential expression (p-values ≤ 0.05) are shown in red when up-regulated (e.g., higher expression in anterior, saddle, flank, or posterior/tail domains for the dorso-ventral axis) or green when down-regulated. Differentially expressed pigmentation genes, homeobox factors and other relevant candidates are shown in grey boxes.

**Fig. S12.**
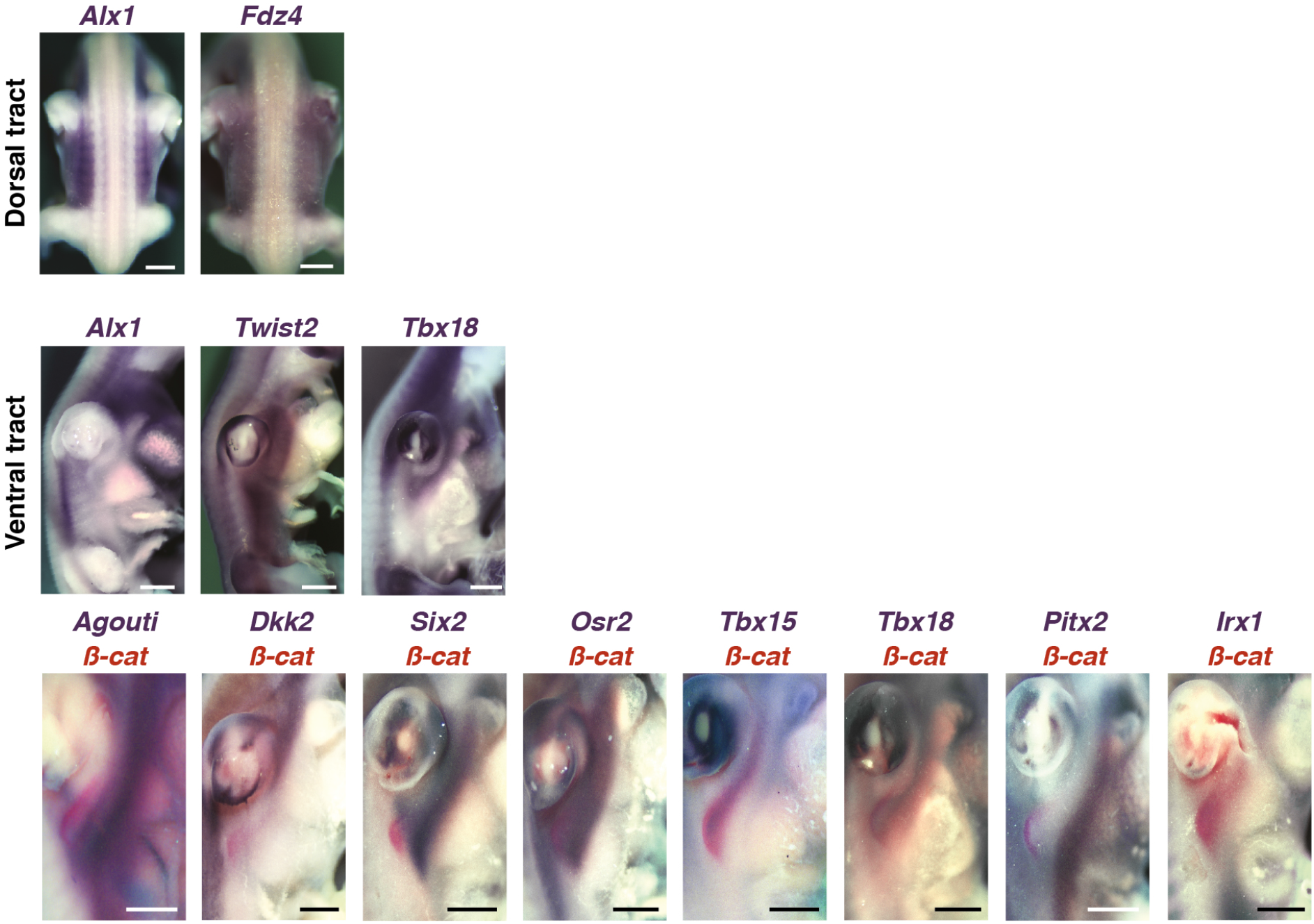
Expression of spatially-restricted candidate genes in the zebra finch embryo. *In situ* hybridizations for genes indicated in purple in the dorsal and ventral regions of zebra finch embryos at stage hh28 confirm quantitative RNA-seq data and qualitatively show they possess spatially-restricted expression patterns (e.g., profiles of *Alx1* and *Twist2* create visible boundaries between breast and ventral domains and *Tbx18* marks the flank domain). The nascent ventral tract was stained using *β-catenin* (in red) to assess the extent of expression domains of candidates encompassing only part of the tract. Scale bars: 500 μm.

**Fig. S13.**
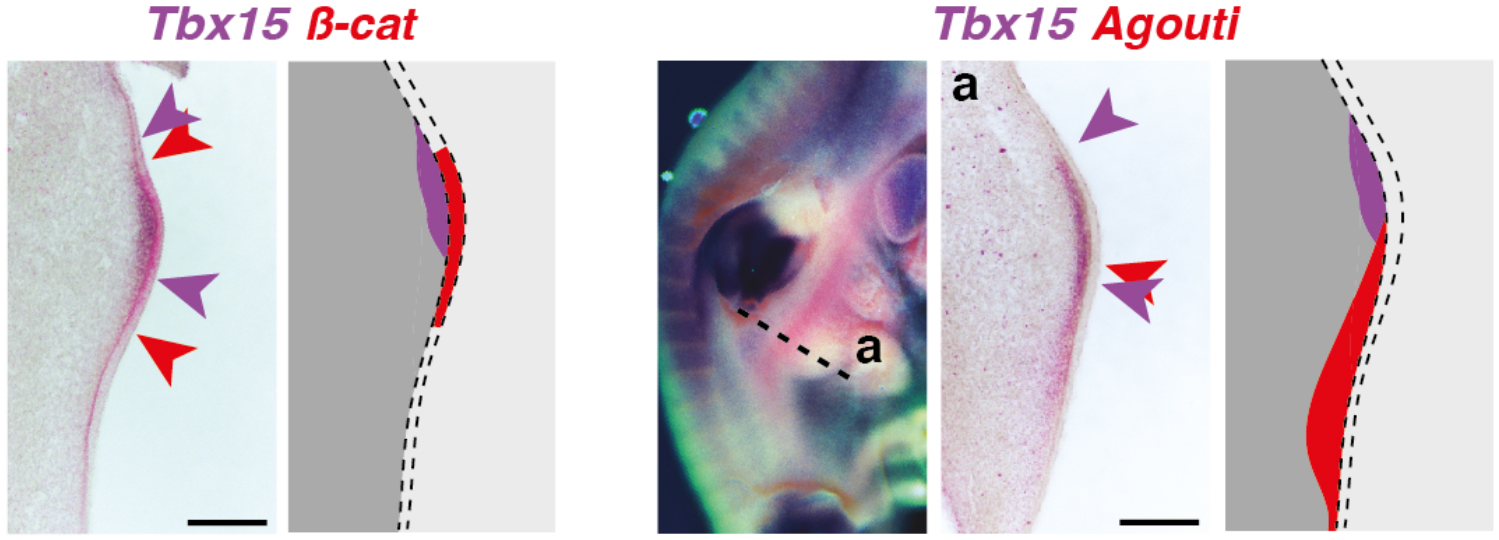
Ventral expression of *Tbx15* and *Agouti* in the zebra finch embryo. Double *in situ* stains for *Tbx15* (in purple) and *β-catenin* or *Agouti* (in red) show that *Tbx15* expression locates in a region directly dorsal to the tract, in a pattern complementary to *Agouti,* precisely marking the location of the future flank-ventral boundary. Scale bars: 100 μm.

**Fig. S14.**
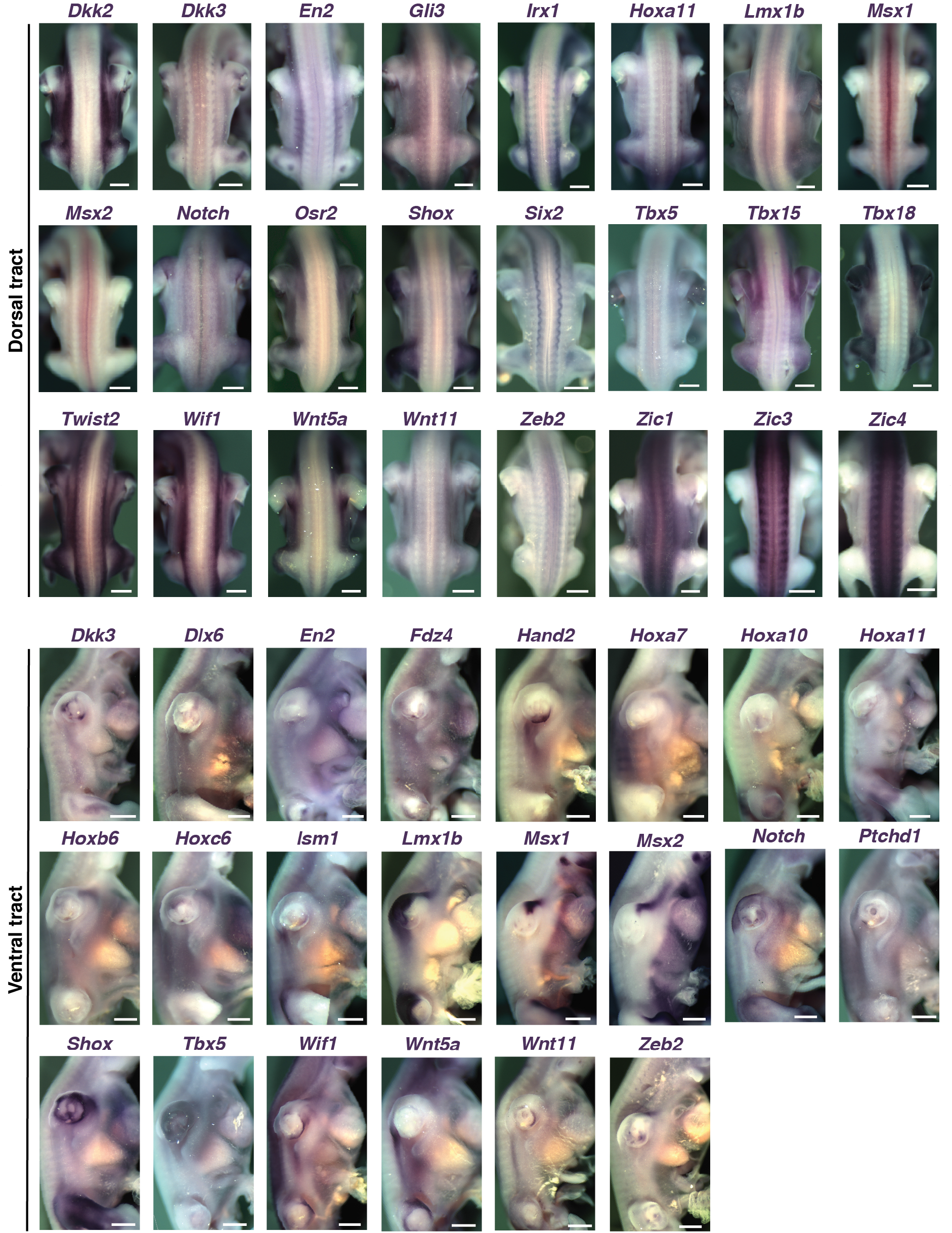
Expression of other candidate genes in the zebra finch embryo. *In situ* hybridizations for genes indicated in purple in the dorsal and ventral regions of zebra finch embryos at stage hh28 confirm quantitative RNA-seq data and qualitatively show they are expressed outside of presumptive tracts and/or throughout tract length with varying widths, thus not spatially correlating with observed boundaries (see Fig. S5). Scale bars: 500 μm.

